# Genomic insights into *Candidatus* Argioplasma dusa, a bacterial symbiont of the wasp spider *Argiope bruennichi*

**DOI:** 10.64898/2026.06.03.729754

**Authors:** Larissa M. Busch, Monica M. Sheffer, Nina Dombrowski, Henrik Krehenwinkel, Stefan Prost, Anja Spang, Gabriele Uhl, Tim Urich, Katharina J. Hoff, Mia M. Bengtsson

**Author notes:** shared first authorship (Larissa M. Busch, Monica M. Sheffer).

## Abstract

Bacterial symbionts were shown to have significant beneficial impacts on the fitness of their host in insects, but little is known on the symbionts of spiders and their interactions with their hosts.

Here, we assembled and investigated the circular 575 kb genome of *Candidatus* Argioplasma dusa, a novel bacterial symbiont of the wasp spider *Argiope bruennichi*. Phylogenomic analysis placed this species within the phylum Tenericutes, in a poorly characterized clade that may represent a new order-level lineage or affiliate with the Mycoplasmatales order. With 559 predicted genes, the genome is relatively small compared to other Tenericutes genomes (on average 969 genes) and has a low GC content of ~24%. While the genome encodes genes for proteins involved in glycolysis and fermentative acetate production, it revealed minimal biosynthetic capabilities with pathways for nucleotide, amino acid and vitamin biosynthesis being absent in *Ca*. Argioplasma dusa. This suggests an intracellular endosymbiotic lifestyle within the spider host. The symbiont was detected in *A. bruennichi* populations across the distribution range of the spider but appears to be absent in certain populations.

## Introduction

Host-adapted bacteria influence the biology of their hosts in many ways, as reproductive parasites, commensals or mutualists (Kaltenpoth *et al*., 2025). Approximately one third of all sequenced bacterial genomes are from those that live in close interactions with eukaryotic hosts (Toft and Andersson, 2010). Depending on their level of host dependency and the intimacy of their interactions, their genomic characteristics differ widely. For instance, obligate intracellular bacterial insect symbionts are often characterized by extremely reduced genomes (Moran and Bennett, 2014) that have evolved independently in multiple bacterial clades from free-living bacterial ancestors with larger genomes (Rogers *et al*., 1985; Ochman, 2005; Moran and Bennett, 2014). The loss of genes in symbionts can be explained by their increasing reliance on gene products of their host organism.

Although endosymbiosis is well-documented in a number of insect taxa and especially well-explored in those groups that exploit challenging nutritional resources such as leafhoppers, cicadas, aphids and some beetles (Bennett, Kwak and Maynard, 2024; Kaltenpoth *et al*., 2025), the currently known host taxa do not reflect the diversity of potential arthropod hosts, thereby limiting our understanding of the range of arthropod-bacterial symbiosis, their functional and evolutionary roles, as well as their genomic features (Mathieson *et al*., 2025).

In particular, spiders are among the less-studied arthropod groups. However, reproductive parasites such as *Wolbachia, Rickettsia, Cardinium* and *Spiroplasma* and additional symbiont species have been discovered in several spider species (Vanthournout and Hendrickx, 2015; Zhang *et al*., 2017, 2018; Sheffer *et al*., 2020; White *et al*., 2020; Perez-Lamarque *et al*., 2022; Nariman *et al*., 2025). Spiders have an obligate predatory lifestyle (Nentwig, 1987), which implies potential transmission of symbionts into spiders from their prey (Kennedy *et al*., 2020), as well as specialized nutritional roles of symbionts within their spider hosts.

A genomic study of two populations (Germany, Estonia) of the wasp spider *Argiope bruennichi* revealed the presence of a previously unknown bacterial endosymbiont (Sheffer *et al*., 2020, 2021) referred to as DUSA (dominant unknown symbiont of *Argiope*). Here, we refer to this symbiont as *Candidatus* Argioplasma dusa. The fact that the symbiont was found in multiple tissues (leg, prosoma, hemolymph, book lungs, ovaries, silk glands, midgut, and fecal pellets; Sheffer *et al*., 2020) demonstrates that it is not part of the microbiome of the digestive tract but likely represents an endosymbiont. DUSA was shown to be affiliated with the *Mollicutes*, the major class in the phylum *Tenericutes*, though the 16S rRNA gene had a remarkably low identity (<85%) to 16S rRNA genes in public databases (Sheffer *et al*., 2020).

Currently known representatives of Mollicutes are small microorganisms (<1µm in diameter) that lack a cell wall (Gasparich, 2010; Razin and Hayflick, 2010). The major genera of the Mollicutes, *Mycoplasma* and *Spiroplasma*, are widespread and comprise mutualists and parasites of many animals and plants (Razin, Yogev and Naot, 1998; Yavlovich, Tarshis and Rottem, 2004; Gasparich, 2010; Razin and Hayflick, 2010; Zhang *et al*., 2018; White *et al*., 2020).

Here, we assembled the full circular genome of *Ca*. A. dusa from metagenome datasets, that were originally used to construct the genome of the host spider *A. bruennichi* (Sheffer *et al*., 2021) and to describe its global expansion range (Krehenwinkel, Rödder and Tautz, 2015). Our genomic and phylogenetic analyses reveal its metabolic potential and phylogenetic placement within the Tenericutes. Furthermore, by investigating metagenomes of geographically separated European and Japanese spider host populations, we were able to shed light into the biogeography of *Ca*. Argioplasma dusa and its population dynamics.

## Materials and Methods

### Genome assembly and annotation

We used previously published (meta)genome sequencing reads that were originally produced for assembling the wasp spider genome (Sheffer *et al*., 2021), but also containing endosymbiont genome reads, to generate the *Ca*. Argioplasma dusa symbiont genome assembly. We assessed three different assembly strategies to separate *A. bruennichi* genome information, *Ca*. A. dusa genome information and contaminant/off-target genetic information (Fig. S1). The strategies and assembly methods are described in detail in the supplementary method S1. In brief, we used PacBio whole genome reads (Sequence Read Archive: SRR11652932; (Sheffer *et al*., 2021)) of a female *A. bruennichi* from Portugal to generate the baseline genome assembly, and Illumina paired-end reads (European Nucleotide Archive: ERR574428; (Krehenwinkel, Rödder and Tautz, 2015)) of several populations for assembly polishing. PacBio reads were trimmed after removing duplicated reads by SeqKit rmdup (v0.12.0; (Shen *et al*., 2016)) using Filtlong (v0.2.0; settings:--min_length 1000 --keep_percent 90; github.com/rrwick/Filtlong) and Illumina reads were trimmed using Trimmomatic (v0.39; settings: ILLUMINACLIP:TruSeq3-PE.fa:2:30:10:2:keepBothReads LEADING:3 TRAILING:3 MINLEN:36; (Bolger, Lohse and Usadel, 2014)).

To create the final selected assembly, we binned the PacBio reads using BlobTools (v1.1.1; (Laetsch and Blaxter, 2017)) and mapping of hits to the UniProt database (release: 2019_02; (The UniProt Consortium, 2019)) by DIAMOND (v0.9.35.136; (Buchfink, Xie and Huson, 2015)). Reads taxonomically classified as “Tenericutes” were selected. These reads were used as the starting assembly for a “map-and-assemble” approach (Wang and Chandler, 2016): all reads were iteratively mapped to the assembly using Minimap2 (v2.17; settings: -ax map-pb; (Li, 2018)) and all mapped reads were re-assembled with Flye (v2.7; settings: --genome-size 1m; (Kolmogorov *et al*., 2019)) until there was no increase of assembly genome size. The resulting assembly was used as a draft assembly, after which all mapping reads were re-assembled using Raven (v1.1.10; (Vaser and Šikić, 2019)) and polished twice with Racon (v1.4.15; (Vaser *et al*., 2017)) using the PacBio reads against the assembly by Minimap2 and five times with Pilon (v1.23; (Walker *et al*., 2014)) using the Illumina reads mapped against the assembly by BWA-MEM (v0.7.17; (Li, 2013)) and sorted with SAMtools (v1.10; (Li *et al*., 2009)). The polished assembly was checked for contaminants using BlobTools, for circularity using Bandage (v0.8.1; (Wick *et al*., 2015)) and assembly metrics were computed with QUAST (v5.0.2; (Gurevich *et al*., 2013)). Additionally, the 16S rRNA gene in the assembly was identified and compared to that of the symbiont identified in the previous *A. bruennichi* microbiome study (Sheffer *et al*., 2020) using NCBI-Blast+ (v2.10.1+; (Camacho *et al*., 2009)). The assembly was submitted under accession CP070274, BioProject: PRJNA629526 and BioSample: SAMN17496637 at the NCBI. Based on the PGAP (Tatusova *et al*., 2016) structural gene annotation obtained from the RefSeq pipeline, completeness of the genome assembly was estimated with BUSCO (v4.1.1;(Seppey, Manni and Zdobnov, 2019)) using the Tenericutes OrthoDB v10 (Kriventseva *et al*., 2019) reference database.

### Phylogenetic analysis

Maximum likelihood phylogenetic reconstructions were performed using 26 single-copy markers from a previously established marker gene set and a backbone of 1,205 bacterial genomes (Moody *et al*., 2022). To identify marker protein homologs, proteins from all genomes of interest were combined and used in an hmmsearch (v3.1b2, settings: hmmsearch --tblout output--domtblout --notextw; (Finn, Clements and Eddy, 2011)) against the COG profiles database (Tatusov *et al*., 2003). Afterwards, the 26 markers were extracted and if more than one hit was assigned to each marker for a single genome, only the best hit was retained based on the best bit-score and e-value. Marker proteins were individually aligned using MAFFT L-INS-i (v7.407, settings: --anysymbol --reorder; (Katoh, 2002)) and trimmed using BMGE (v1.12, settings: -t AA -m BLOSUM30 -b 2 -h 0.55; (Criscuolo and Gribaldo, 2010)). The trimmed alignments were concatenated using catfasta2phyml (github.com/nylander/catfasta2phyml) and a phylogenetic tree was generated using IQ-TREE (v2.1.2; settings: -m LG+F+R -bb 1000 -alrt 1000; (Minh *et al*., 2020)). This tree was used to down sample the taxon selection to 479 bacterial genomes (Table S1; Figure S1). From this taxon set, the 26 marker proteins were extracted, aligned and trimmed using MAFFT L-INS-i and BMGE as described above. Alignments were concatenated and a phylogenetic tree was generated using IQ-TREE (settings: - m LG+C60+F+R -bb 1000 -alrt 1000).

SlowFaster (v1; (Kostka *et al*., 2008)) was used with default settings on the concatenated alignment to remove fast-evolving sites. Additionally, heterogeneous sites were removed using alignment_pruner.pl (settings: --chi2_prune, github.com/novigit/davinciCode/blob/master/perl). For both approaches, sites were removed in a step-wise manner removing 10, 20, 30 and 40% of sites (Figures S2-9). Afterwards, phylogenetic trees were generated using IQ-TREE (settings: -m LG+C20+F+R -bb 1000 -alrt 1000).

For further functional annotation, we analyzed the *Ca*. A. dusa symbiont genome and 478 bacterial genomes. Rapid gene calling was performed using Prokka (Seemann, 2014), with genetic code 11 applied to all genomes except Mycoplasma, for which genetic code 4 was used. Afterwards, the generated proteins were compared against several databases (supplementary methods S2).

### Population genetic analysis

SNP analysis to investigate the population structure of *Ca*. A. dusa was conducted using Illumina reads obtained from *A. bruennichi* populations in Japan (five specimen data sets), Sweden, the Baltic region, Portugal, Italy (European Nucleotide Archive: ERR574428; (Krehenwinkel, Rödder and Tautz, 2015), the Azores and Madeira. The population specific reads were mapped against the *Ca*. A. dusa genome by BWA-MEM (v0.7.17; (Li, 2013)). SNP calling was then performed using SAMtools and BCFtools (v1.10; (Danecek *et al*., 2021)) assuming a ploidy of 1. The top 90 % quality sites identified in all populations were analyzed by principal component analysis (PCA) and hierarchical clustering in R (v4.1.2) using the packages tidyverse (v2.0.0; (Wickham *et al*., 2019)), FactoMineR (v2.8; (Lê, Josse and Husson, 2008)) and factoextra (v1.0.7) to characterize the *Ca*. A. dusa population differences according to the variant calls.

## Results

### *Ca*. Argioplasma dusa genome characteristics suggesting genome reduction

The genome assembly of *Ca*. A. dusa (accession CP070274) using a metagenomics approach (Supplementary Methods S1), resulted in a set of 18 candidate assemblies. We selected the assembly with the largest genome size, 575,551 bp, present in one circular contig as representative assembly for the genome of *Ca*. A. dusa genome (Table 1). The single 16S rRNA gene detected on this contig is 99.8% identical to the previously reported full-length 16S rRNA gene sequence of *Ca*. A. dusa in the *A. bruennichi* microbiome (Sheffer *et al*., 2020). The genome contains 526 protein coding sequences (CDS) in addition to 27 tRNA genes and one of each for 5S, 16S and 23S rRNA gene, respectively, as well as three ncRNAs.

**Table 1:** Assembly statistics. Assembly metrics were computed with QUAST v5.0.2 (Gurevich *et al*., 2013) and DUSA 16S rRNA (Sheffer *et al*., 2019) gene sequence identity was analyzed with Blast+ (Camacho *et al*., 2009). CDS Annotation done with NCBI Prokaryotic Genome Annotation Pipeline (PGAP) (Tatusova *et al*., 2016).

| Assembly statistics |  |
| --- | --- |
| # contigs | 1 |
| Assembly size [bp] | 575,551 |
| N50 [bp] | 575,551 |
| GC (%) | 24.12 |
| 16S rRNA gene identity | 99.8% |
| # genes | 559 |
| Complete BUSCOs | 82.7% |
| Fragmented BUSCOs | 1.3% |
| Missing BUSCOs | 16.0% |

The previous analysis of the 16S rRNA gene sequence placed the endosymbiont into the clade Tenericutes (nov. Mycoplasmatota) (Sheffer *et al*., 2020). Comparison with 102 Tenericutes genomes in the RefSeq database, which range from 0.56-1.97 Mb (mean: 1.03) and encode for 560-4,727 genes (mean: 969), revealed that the genome of *Ca*. A. dusa at 0.58 Mb was among the smallest Tenericutes characterised to date. This could reflect a reduced genomic complexity often observed for endosymbionts (Román-Escrivá *et al*., 2025). In agreement with the previous observation that many (endo)symbiont genomes have an AT bias with a consequently low GC content (McCutcheon, McDonald and Moran, 2009; Román-Escrivá *et al*., 2025), the *Ca*. A. dusa genome had a low GC content (24.1%), even lower than that of most Tenericutes reference genomes (range 23.1-40.0%; mean 27.9%) and also lower than the GC content of the *A. bruennichi* genome (29.3%; (Sheffer *et al*., 2021)). *Ca*. A. dusa encoded 82.7% (n=124/150 genes) of the complete Tenericutes OrthoDB v10 marker genes (Kriventseva *et al*., 2019), while 1.3% of genes (n=2) were fragmented and 16% were missing (n=24). None of the 24 missing Tenericutes marker genes (Supplementary Data Table S2) were identified in any of the alternative genome assemblies, indicating that these genes are likely missing in the *Ca*. A. dusa genome.

### *Ca*. A. dusa falls into a deep-branching Tenericutes clade

Phylogenetic analyses of *Ca*. A. dusa using 26 marker genes revealed its putative placement within the class Mollicutes in the phylum Tenericutes. Together with three other taxa, *Ca*. A. dusa may either be affilated with the Mycoplasmatales order or form a separate order-level lineage sister to Mycoplasmatales or Mycoplasmatales and Entomoplasmatales clades combined (Figure 1 & S1-9). Specifically, while the full alignment recovered a placement between Mycoplasmatales and Mycoplasmatales/Entomoplasmatales, upon filtering of the 30-40% most compositionally or fast-evolving sites, this clade emerged basal to all Mycoplasmatales and Entomoplasmatales (Fig. 1A). The closest sequenced relatives to *Ca* A. dusa are characterized by similarly small genome sizes, and were found in a snail (host: *Biomphalaria glabrata*, symbiont genome size: 0.65 Mb, accession number: GCA_001484045; (Adema *et al*., 2017)) and a glass sponge collected from the Scotia Shelf (host: *Vazella pourtalesii*, symbiont genome size: 0.67 Mb & 0.68 Mb,GCA_014238525 & GCA_014238225; (Bayer *et al*., 2020)).

**Figure 1:**
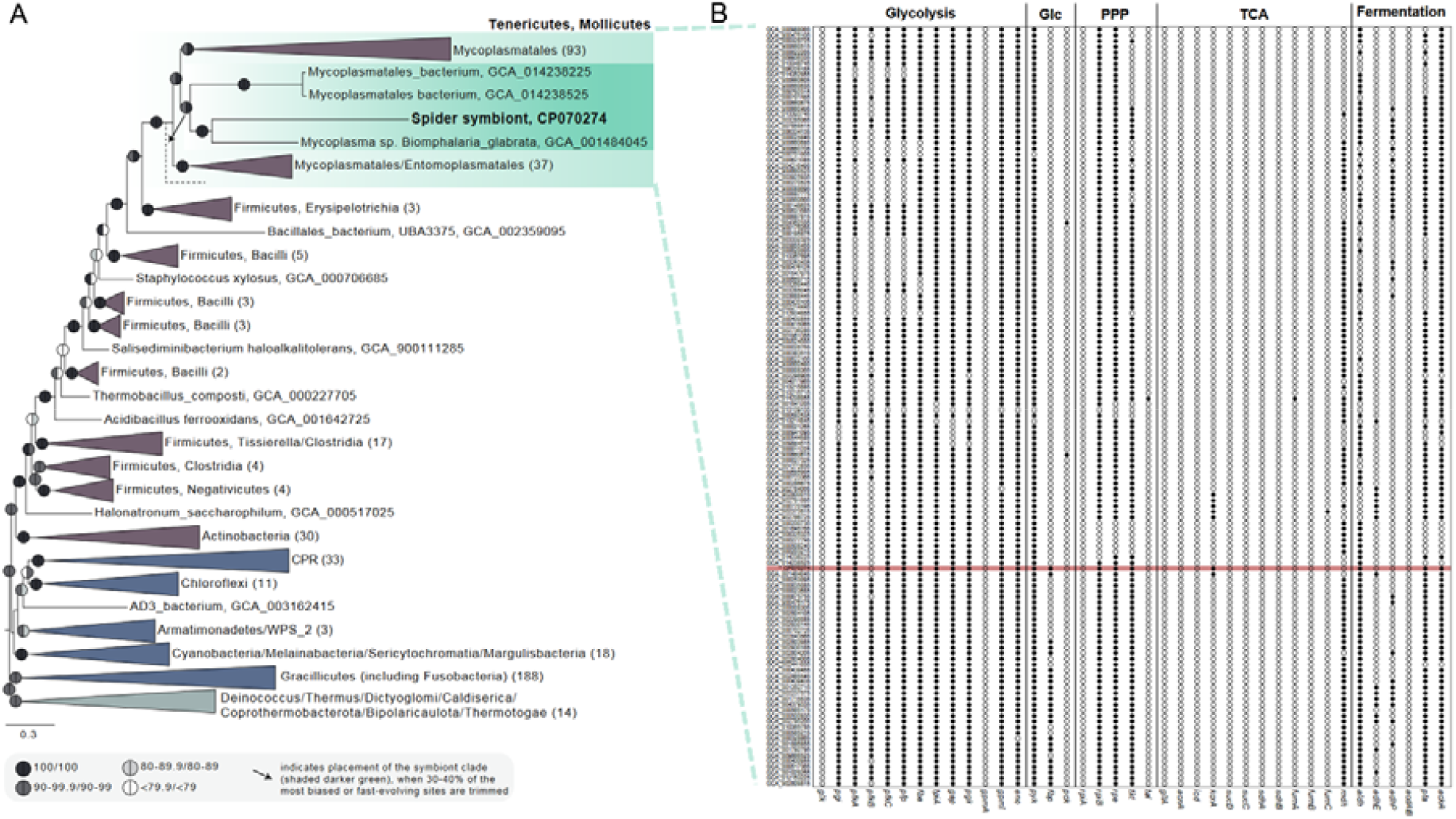
Phylogenetic placement of *Ca*. A. dusa (A). Comparison of predicted C-metabolism of Mycoplasmatales (B). Gene content of *Ca*. A. dusa is highlighted in red. Arrow points to the alternative position obtained when removing 30-40% of the most compositionally or fast-evolving biased sites (see Supplementary Figure 1 and 2).

The long branches separating those Tenericutes from each other and their next relatives, may indicate that these organisms have experienced relatively fast evolutionary rates due to their symbiotic lifestyle as commonly observed in recent insect symbionts (O’Fallon, 2008; Bennett *et al*., 2014; Gerth *et al*., 2021; Wang *et al*., 2026). Conversely, the phylogenetic diversity of putative hosts indicates that this Tenericutes clade is likely currently under-sampled and may comprise additional members infecting a broad range of eukaryotic host families.

### Functional gene content and metabolic potential

Similar to other Tenericutes genomes (Pollack, Williams and McElhaney, 1997), the genome of *Ca*. A. dusa encodes genes for complete glycolysis pathway, while lacking genes for the oxidative citric acid cycle (Fig. 2A & Fig S11, Table S3). Instead, acetyl-CoA, produced from pyruvate oxidoreductase (*porA*) after glycolysis, is likely transformed to acetate in a short fermentative pathway *via* phosphate acetyltransferase (*pta*) and acetate kinase (*ackA*, Fig.2) yielding ATP *via* substrate level phosphorylation. No genes encoding proteins of aerobic or anaerobic respiratory chains were detected. Diverse simple carbohydrates, such as fructose, glucose, mannose, galactose, N-acetyl-glucosamine are predicted to be fueled into the glycolysis pathway based on the detected PTS system components and downstream enzymes (*pgi, pfkA, pfkC, ptp, fba, tpiA, gap, pgk, gpmI, eno, pyk*) (Fig. 1B & 2A, Table S3). Differing from most Tenericutes species, *Ca*. A. dusa also encodes genes involved in gluconeogenesis (*fbp*; Figure 1B). Given this, *Ca*. A. dusa might provide acetate and/or monosaccharides to its spider host. No genes encoding enzymes of fatty acid biosynthesis are found; however, the genes encoding enzymes of the terminal steps of phospholipid synthesis are identified (*plsC, pgsA, CIs*), indicating a dependency on host fatty acids commonly observed in genome reduced endosymbionts (McCutcheon *et al*., 2024). The small *Ca*. A. dusa genome encodes very few enzymes for amino acid biosynthesis, again indicating metabolic dependency on the spider host. While the nucleic acid metabolism is predicted to be similar to other Tenericutes, we did not find any alternative vitamin biosynthesis pathways (Table S3).

**Figure 2:**
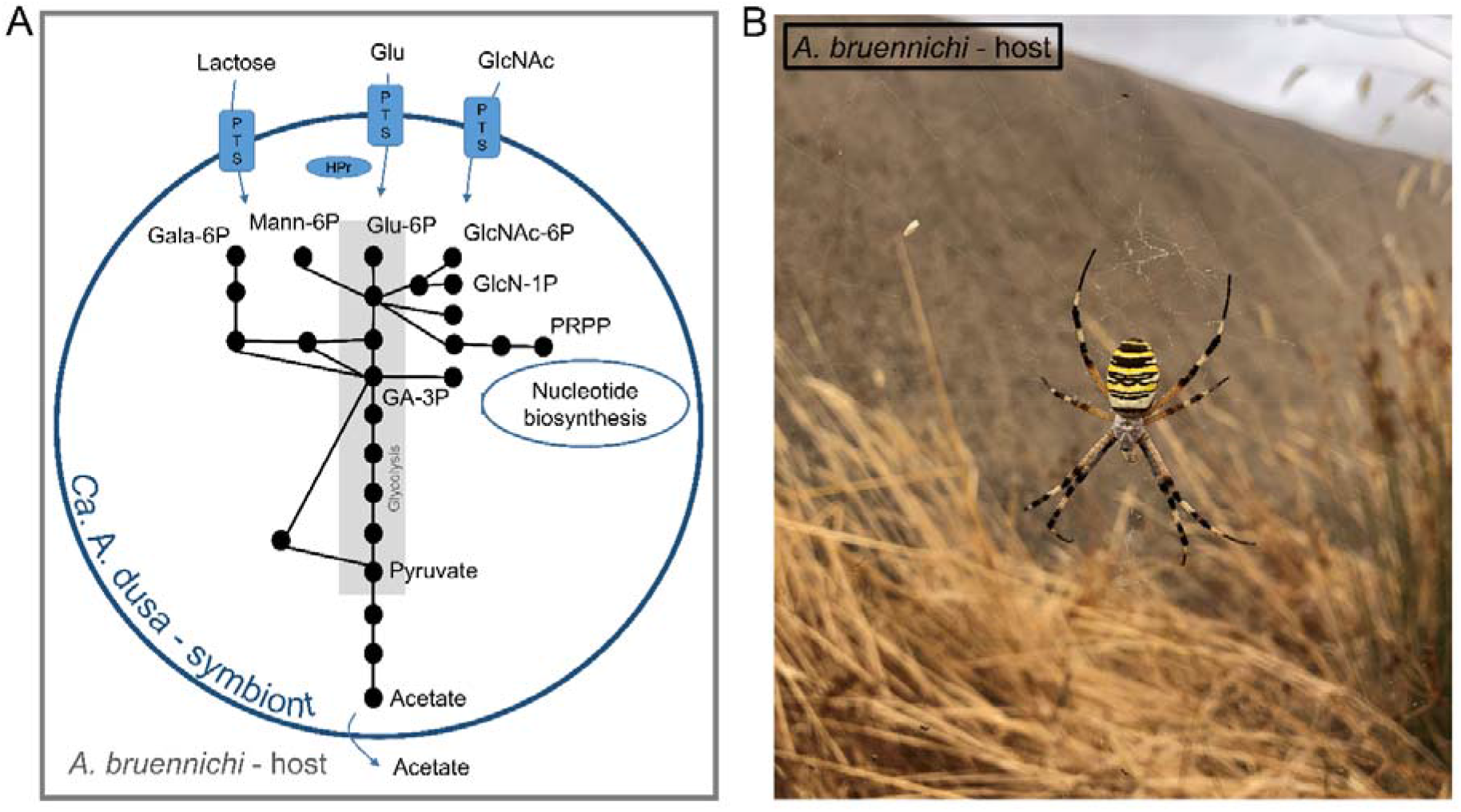
Model of the central C-metabolism of *Ca*. A. dusa. (A) Based on the functional annotation of *Ca*. A. dusa genes, the metabolic potential is depicted. Sugars are likely provided by the *A. bruennichi* host, whereas the fermentative product acetate is provided by the symbiont and further metabolized by the host. Glu: Glucose. GlcNAc: N-Acetylglucosamine. Gala-6P: Galactose-6-phosphate. Mann-6P: Mannose-6-phosphate. Glu-6P: Glucose-6-phosphate. GlcNAc-6P: N-Acetylglucosamine-6-phosphate. GlcN-1P: Glucosamine-1-phosphate. PRPP: Phosphoribosyl pyrophosphate. GA-3P: Glyceraldehyde-3P. Detailed annotation is given in Fig. S2. (B) Photo of the host organism *A. bruennichi* taken in 2019 of a specimen in Portugal.

### *Ca*. A. dusa populations differ in regard to their geographic distribution

The thermophilic wasp spider *A. bruennichi* (Fig. 2B) has a broad global distribution and is currently undergoing a natural poleward range expansion partly caused by climate change (Krehenwinkel and Tautz, 2013; Krehenwinkel and Pekar, 2015; Krehenwinkel, Rödder and Tautz, 2015; Krehenwinkel *et al*., 2016; Wawer *et al*., 2017; Wawer and Wytwer, 2020; Wolz *et al*., 2020; Sheffer *et al*., 2026). Therefore, we asked if all populations carry the *Ca*. A. dusa symbiont and if there are geographic differences in the *Ca*. A. dusa populations as the carriage of *Ca*. A. dusa has only been demonstrated for German and Estonian *A. bruennichi* populations (Sheffer *et al*., 2020).To shed light on the presence and genome variation of *Ca*. A. dusa in different populations of the spider host, we performed single nucleotide variant (SNV) analysis of seven geographically distinct populations. Mapping of Illumina reads from such distinct *A. bruennichi* populations to the *Ca*. A. dusa genome revealed a separation of *Ca*. A. dusa genomes according to theier geographic origin. Of note, not all investigated populations harbored the symbiont; specifically, it appears to be absent from three of the five analyzed Japanese populations (Table S4).

Principal component analysis (PCA) and hierarchical clustering of Japanese, European and Atlantic Ocean populations according to the SNV calls present in all analyzed populations shows that the geographic distribution is reflected by the genetic variance of *Ca*. A. dusa (Fig. 3A & B) suggesting different *Ca*. A. dusa lineages. The divergence of microbial genomes also reflects the Palearctic-wide phylogeographic divergence pattern of the host spider (Krehenwinkel *et al*., 2016), which further supports the assumption of a host-symbiont co-evolutionary process.

**Figure 3:**
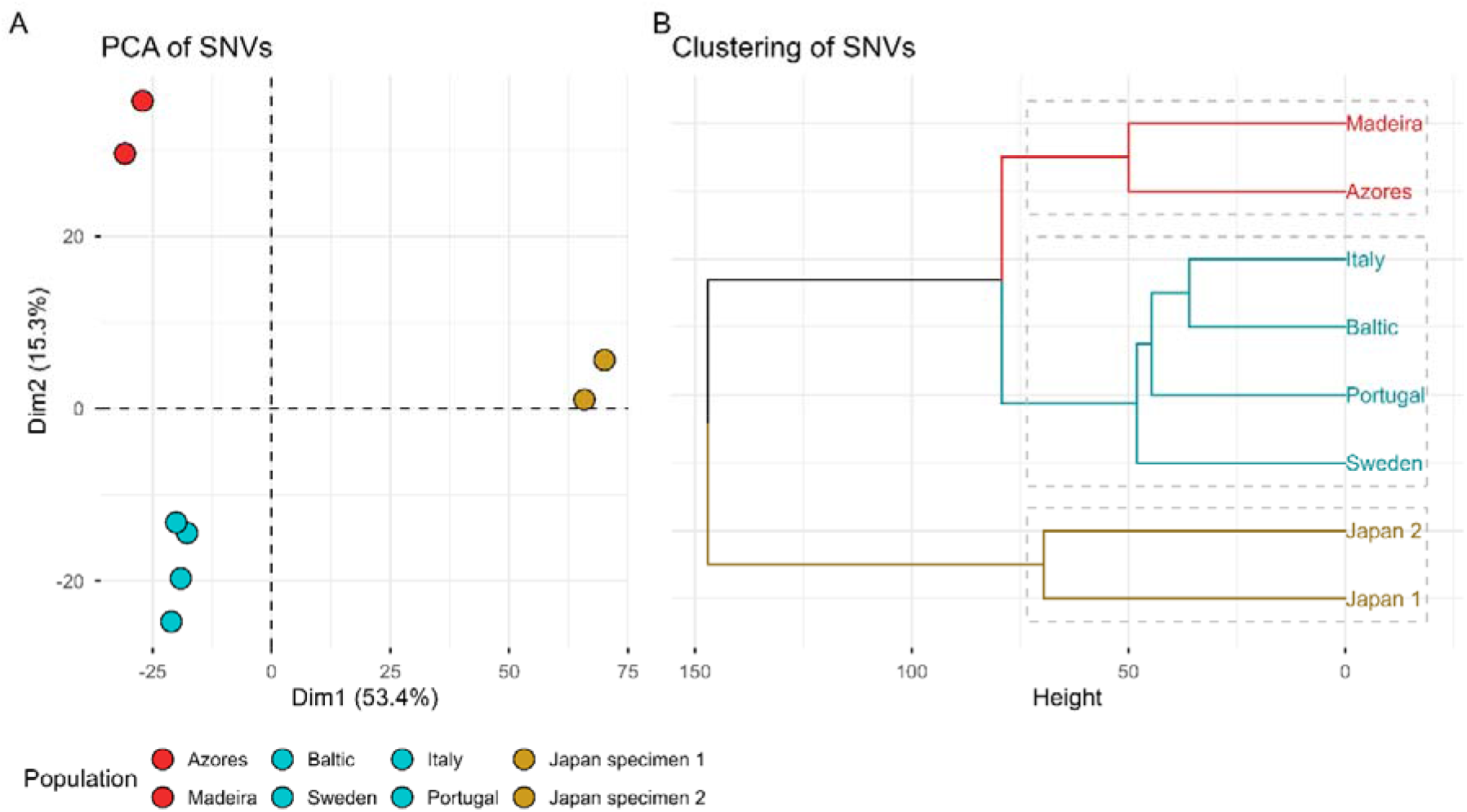
Genome SNV analysis of *Ca*. A. dusa from different populations of *A. bruennichi*. Illumina reads generated for *A. bruennichi* populations/specimens from Japan, Europe (Baltic region, Sweden, Italy, Portugal) and Atlantic islands (Azores, Madeira) were mapped to the *Ca*. A. dusa genome.Corresponding *Ca*. A. dusa population SNV calls were visualized as principal component analysis (A) and hierarchically clustered (B).

## Discussion

A proposed symbiont of Tenericutes clade was first detected in *A. bruennichi* spider populations from Germany and Estonia *via* 16S rRNA gene amplicon sequencing (Sheffer *et al*., 2020). To study this symbiont in detail, we assembled and annotated the whole genome of this organism using a metagenomics approach from shotgun sequencing data of whole *A. bruennichi* tissues (Sheffer *et al*., 2021). This approach highlights the potential of reanalysis of available metagenomics data to discover bacteria that occur in natural populations of diverse hosts and provide first insights into the possible role they may play in their host biology and (co-)evolutionary history.

Our current assembly and phylogenetic analyses of the symbiont demonstrates that this organism is only distantly related to other known symbionts of the orders Mycoplasmatales and Entomoplasmatales within the Mollicutes class in the phylum Tenericutes. The closest currently known related genomes of *Ca*. Argioplasma dusa were recovered from such diverse hosts as the freshwater snail *Biomphalaria glabrata* (Adema *et al*., 2017) and a marine glass sponge *Vazella pourtalesii* (Bayer *et al*., 2020). That the symbiont bacteria are associated to hosts, which are scattered across the tree of life and occur in very different habitats, may be a consequence of missing observations from other host species and habitats. However, a wider search of the 16S rRNA gene sequence of *Ca*. A. dusa against the NCBI SRA (sequence read archive) did not recover any closely related sequences from other spiders or insects (data not shown) and *Ca*. A. dusa was also not identified in a range of closely related Argiope species (H. Krehenwinkel, personal observation).

This raises questions about the origin and dynamics of the symbiotic interaction with the wasp spider. Reanalysis of metagenomics sequencing data of several *A. bruennichi* populations indicated that only two out of five Japanese populations harbor *Ca*. A. dusa, which might suggest that it is not an obligate, primary endosymbiont. The Japanese population is likely the most basal population of the studied spider populations as eastern Asia as a major glacial refugium (Krehenwinkel *et al*., 2016). Consequently, it remains to be determined whether the symbiosis was ancestral to this spider species with subsequent loss in some populations or was acquired independently in some populations later on. However, reads mapping to the symbiont genome in samples from *A. bruennichi* populations varied across several SNVs, in a pattern consistent with geographic distribution and genetic variation of the spider host (Wawer *et al*., 2017), indicating a possible co-evolution of the spider host and the *Ca*. A. dusa symbiont. Given the recent geographic range expansion of the host, this suggests that symbiont evolution may be rapid. In line with other arthropod-symbiont systems (Goodacre and Martin, 2012), we hypothesize that the symbiont may have a potential role in phenotypic traits linked to geographic expansion, such as dispersal, cold tolerance, or reproduction. Indeed, previous work has shown that evolutionary diversification and adaptation of host species is associated with carrying of symbionts (Cornwallis *et al*., 2023; Bennett, Kwak and Maynard, 2024). Analysis of the metabolic potential of *Ca*. A. dusa revealed that it potentially provides acetate and monosaccharides to the spider host, which could be nutritionally beneficial or act as anti-freeze substances for overwintering juvenile *A. bruennichi*. Prospective experimental work should focus on testing hypotheses on the potential functional role of *Ca*. A. dusa in its host. Comparative genomics has provided insights into reductive genome evolution and associated loss in functional abilities of obligate endosymbionts (Moya *et al*., 2008; Moran and Bennett, 2014).

Based on the previous discovery of 16S rRNA genes of *Ca*. A. dusa in all host tissues studied (legs, prosoma, ovaries, midgut, silk glands, hemolymph and fecal pellets) as well as in newly hatched spiderlings, indicates that the symbiont is located within the host body and is maternally transmitted via eggs (Sheffer *et al*., 2020). This points towards a systemic distribution of the symbiont within the host and a potential for vertical transmission to the offspring. However, we cannot distinguish whether the symbiont is intracellular, or otherwise located within the host tissue or present in the hemolymph. The small genome size and level of genome reduction, however, are consistent with *Ca*. A. dusa potentially being an obligate intracellular symbiont, dependent on its host for basic metabolic functions (Moran and Bennett, 2014). Our understanding of the interaction between symbiont and host, including any beneficial or antagonistic effects, is preliminary. The genomic analyses suggest that *Ca*. A. dusa can likely metabolize simple sugars and N-acetyl-glucosamine resulting in acetate as an end product. Given this, *Ca*. A. dusa might provide acetate and/or monosaccharides to its spider host and could be assimilated into the central carbon metabolism, as shown for other arthropod-bacterial symbioses (Ankrah, Chouaia and Douglas, 2018) - however this is highly speculative and requires further investigations. Metabolic interactions between fermentative bacteria producing short-chain fatty acids, such as acetate, propionate or butyrate and their animal host cells are widespread (McFall-Ngai *et al*., 2013).

The absence of biosynthetic pathways for vitamins, amino acids or specialized enzymes for digesting typical dietary components of the spider (insect biomass), do not allow us to infer any typical mutualistic interactions between host and symbiont.

Altogether, the evolutionary history and symbiotic function of *Ca*. A. dusa in the spider *A. bruennichi* remains enigmatic. We encourage future studies to widen the search for *Ca*. A. dusa and related organisms in other potential hosts to explore the role of *Ca*. A. dusa in symbiosis. A logical next step to investigate the specific role for *A. bruennichi* would be to explore whether the symbiont is intercellular or extracellular in the tissues of *A. bruennichi* and to carry out experimental studies on populations with and without symbiont to shed further light onto the functional characteristics of the symbiosis between *A. bruennichi* and *Ca*. A. dusa.

### Description of *Candidatus* Argioplasma dusa

We generated the symbiont genome assembly based on whole genome sequencing reads of its host *Argiope bruennichi*. The symbiont was first detected in *A. bruennichi* populations from Germany and Estonia *via* 16S rRNA gene amplicon sequencing (Sheffer *et al*., 2020). Phylogenetics revealed that the genome is not closely related to known Tenericutes clade genomes. Detection of the symbiont in newly hatched spiderlings suggests vertical transmission to the offspring (Sheffer *et al*., 2020). Distinctive features include strong wasp spider host association, low GC content and a unique 16S rDNA gene sequence (Locus tag JTY60_02675; ncbi.nlm.nih.gov/datasets/gene/GCA_051094275.1/?gene_type=rRNA)

We therefore propose to treat this symbiont as a new candidate bacterial species, under the name *Candidatus* Argioplasma dusa.

‘*Candidatus* Argioplasma’ (Argio.plas.ma; *Argio* – referring to the host *Argiope bruennichi*; *plasma* – referring to class Mycoplasmatales)

*‘Candidatus* Argioplasma dusa’ (dusa; dusa - previous designation as “Dominant Unknown Symbiont of *Argiope bruennichi”* (DUSA), as well as the word **“dúša”**, meaning spirit or soul in Slavic languages). Type material is the genome assembly designated as CP070274.1 (NCBI GenBank) and GCF_051094275.1 / GCA_051094275.1 (NCBI Genome) representing ‘*Candidatus* Argioplasma dusa’.

## Acknowledgments

This work was supported by the DFG (Deutsche Forschungsgemeinschaft) for the research training group (RTG 2010) to G. U., the Swedish Research Council (VR starting grant 2016-03559 to A. S.), the NWO-I foundation of the Netherlands Organisation for Scientific Research (WISE fellowship to A. S.) and the European Research Council (ERC) under the European Union’s Horizon 2020 research and innovation programme (grant agreement ERC Starting grant No. 947317 (ASymbEL) to A. S.).

## Availability of data and materials

- Whole genome *A. bruennichi* PacBio reads (Sequence Read Archive: SRR11652932)
- Whole genome *A. bruennichi* Illumina paired-end reads (European Nucleotide Archive: ERR574428)
- Assembly and annotation: accession CP070274 BioProject: PRJNA629526 BioSample: SAMN17496637 Assembly: GCF_051094275.1 / GCA_051094275.1
- Additional data is available from the corresponding author on request

## Notes

### Competing Interest Statement

The authors have declared no competing interest.

## References

Adema, C.M. et al. (2017) “Whole genome analysis of a schistosomiasis-transmitting freshwater snail,” Nature Communications, 8(1), p. 15451. Available at: 10.1038/ncomms15451.

Ankrah, N.Y.D., Chouaia, B. and Douglas, A.E. (2018) “The Cost of Metabolic Interactions in Symbioses between Insects and Bacteria with Reduced Genomes,” mBio. Edited by A. Wilson and E.G. Ruby, 9(5), pp. e01433–18. Available at: 10.1128/mBio.01433-18.

Bayer, K. et al. (2020) “Microbial Strategies for Survival in the Glass Sponge Vazella pourtalesii,” mSystems. Edited by S. Bordenstein, 5(4), pp. e00473–20. Available at: 10.1128/mSystems.00473-20.

Bennett, G.M. et al. (2014) “Differential Genome Evolution Between Companion Symbionts in an Insect-Bacterial Symbiosis,” mBio. Edited by M.J. McFall-Ngai, 5(5), pp. e01697–14. Available at: 10.1128/mBio.01697-14.

Bennett, G.M., Kwak, Y. and Maynard, R. (2024) “Endosymbioses Have Shaped the Evolution of Biological Diversity and Complexity Time and Time Again,” Genome Biology and Evolution. Edited by A. Eyre-Walker, 16(6), p. evae112. Available at: 10.1093/gbe/evae112.

Bolger, A.M., Lohse, M. and Usadel, B. (2014) “Trimmomatic: a flexible trimmer for Illumina sequence data,” Bioinformatics, 30(15), pp. 2114–2120. Available at: 10.1093/bioinformatics/btu170.

Buchfink, B., Xie, C. and Huson, D.H. (2015) “Fast and sensitive protein alignment using DIAMOND,” Nature Methods, 12(1), pp. 59–60. Available at: 10.1038/nmeth.3176.

Camacho, C. et al. (2009) “BLAST+: architecture and applications,” BMC Bioinformatics, 10(1), p. 421. Available at: 10.1186/1471-2105-10-421.

Cornwallis, C.K. et al. (2023) “Symbioses shape feeding niches and diversification across insects,” Nature Ecology & Evolution, 7(7), pp. 1022–1044. Available at: 10.1038/s41559-023-02058-0.

Criscuolo, A. and Gribaldo, S. (2010) “BMGE (Block Mapping and Gathering with Entropy): a new software for selection of phylogenetic informative regions from multiple sequence alignments,” BMC Evolutionary Biology, 10(1), p. 210. Available at: 10.1186/1471-2148-10-210.

Danecek, P. et al. (2021) “Twelve years of SAMtools and BCFtools,” GigaScience, 10(2), p. giab008. Available at: 10.1093/gigascience/giab008.

Finn, R.D., Clements, J. and Eddy, S.R. (2011) “HMMER web server: interactive sequence similarity searching,” Nucleic Acids Research, 39(suppl), pp. W29–W37. Available at: 10.1093/nar/gkr367.

Gasparich, G.E. (2010) “Spiroplasmas and phytoplasmas: Microbes associated with plant hosts,” Biologicals, 38(2), pp. 193–203. Available at: 10.1016/j.biologicals.2009.11.007.

Gerth, M. et al. (2021) “Rapid molecular evolution of Spiroplasma symbionts of Drosophila,” Microbial Genomics, 7(2). Available at: 10.1099/mgen.0.000503.

Goodacre, S.L. and Martin, O.Y. (2012) “Modification of Insect and Arachnid Behaviours by Vertically Transmitted Endosymbionts: Infections as Drivers of Behavioural Change and Evolutionary Novelty,” Insects, 3(1), pp. 246–261. Available at: 10.3390/insects3010246.

Gurevich, A. et al. (2013) “QUAST: quality assessment tool for genome assemblies,” Bioinformatics, 29(8), pp. 1072–1075. Available at: 10.1093/bioinformatics/btt086.

Kaltenpoth, M. et al. (2025) “Origin and function of beneficial bacterial symbioses in insects,” Nature Reviews Microbiology, 23(9), pp. 551–567. Available at: 10.1038/s41579-025-01164-z.

Katoh, K. (2002) “MAFFT: a novel method for rapid multiple sequence alignment based on fast Fourier transform,” Nucleic Acids Research, 30(14), pp. 3059–3066. Available at: 10.1093/nar/gkf436.

Kennedy, S.R. et al. (2020) “Are you what you eat? A highly transient and prey□influenced gut microbiome in the grey house spider Badumna longinqua,” Molecular Ecology, 29(5), pp. 1001– 1015. Available at: 10.1111/mec.15370.

Kolmogorov, M. et al. (2019) “Assembly of long, error-prone reads using repeat graphs,” Nature Biotechnology, 37(5), pp. 540–546. Available at: 10.1038/s41587-019-0072-8.

Kostka, M. et al. (2008) “SlowFaster, a user-friendly program for slow-fast analysis and its application on phylogeny of Blastocystis,” BMC Bioinformatics, 9(1), p. 341. Available at: 10.1186/1471-2105-9-341.

Krehenwinkel, H. et al. (2016) “A phylogeographical survey of a highly dispersive spider reveals eastern Asia as a major glacial refugium for Palaearctic fauna,” Journal of Biogeography, 43(8), pp. 1583–1594. Available at: 10.1111/jbi.12742.

Krehenwinkel, H. and Pekar, S. (2015) “An Analysis of Factors Affecting Genotyping Success from Museum Specimens Reveals an Increase of Genetic and Morphological Variation during a Historical Range Expansion of a European Spider,” PLOS ONE. Edited by M. Kuntner, 10(8), p. e0136337. Available at: 10.1371/journal.pone.0136337.

Krehenwinkel, H., Rödder, D. and Tautz, D. (2015) “Eco□genomic analysis of the poleward range expansion of the wasp spider A rgiope bruennichi shows rapid adaptation and genomic admixture,” Global Change Biology, 21(12), pp. 4320–4332. Available at: 10.1111/gcb.13042.

Krehenwinkel, H. and Tautz, D. (2013) “Northern range expansion of E uropean populations of the wasp spider A rgiope bruennichi is associated with global warming–correlated genetic admixture and population□specific temperature adaptations,” Molecular Ecology, 22(8), pp. 2232–2248. Available at: 10.1111/mec.12223.

Kriventseva, E.V. et al. (2019) “OrthoDB v10: sampling the diversity of animal, plant, fungal, protist, bacterial and viral genomes for evolutionary and functional annotations of orthologs,” Nucleic Acids Research, 47(D1), pp. D807–D811. Available at: 10.1093/nar/gky1053.

Laetsch, D.R. and Blaxter, M.L. (2017) “BlobTools: Interrogation of genome assemblies,” F1000Research, 6, p. 1287. Available at: 10.12688/f1000research.12232.1.

Lê, S., Josse, J. and Husson, F. (2008) “FactoMineR□: An R Package for Multivariate Analysis,” Journal of Statistical Software, 25(1). Available at: 10.18637/jss.v025.i01.

Li, H. et al. (2009) “The Sequence Alignment/Map format and SAMtools,” Bioinformatics, 25(16), pp. 2078–2079. Available at: 10.1093/bioinformatics/btp352.

Li, H. (2013) “Aligning sequence reads, clone sequences and assembly contigs with BWA-MEM.” arXiv. Available at: 10.48550/ARXIV.1303.3997.

Li, H. (2018) “Minimap2: pairwise alignment for nucleotide sequences,” Bioinformatics. Edited by I. Birol, 34(18), pp. 3094–3100. Available at: 10.1093/bioinformatics/bty191.

Mathieson, O.L. et al. (2025) “The ecology, evolution, and physiology of Cardinium□: a widespread heritable endosymbiont of invertebrates,” FEMS Microbiology Reviews, 49, p. fuaf031. Available at: 10.1093/femsre/fuaf031.

McCutcheon, J.P. et al. (2024) “How do bacterial endosymbionts work with so few genes?,” PLOS Biology. Edited by T.A. Richards, 22(4), p. e3002577. Available at: 10.1371/journal.pbio.3002577.

McCutcheon, J.P., McDonald, B.R. and Moran, N.A. (2009) “Origin of an Alternative Genetic Code in the Extremely Small and GC–Rich Genome of a Bacterial Symbiont,” PLoS Genetics. Edited by I. Matic, 5(7), p. e1000565. Available at: 10.1371/journal.pgen.1000565.

McFall-Ngai, M. et al. (2013) “Animals in a bacterial world, a new imperative for the life sciences,” Proceedings of the National Academy of Sciences, 110(9), pp. 3229–3236. Available at: 10.1073/pnas.1218525110.

Minh, B.Q. et al. (2020) “IQ-TREE 2: New Models and Efficient Methods for Phylogenetic Inference in the Genomic Era,” Molecular Biology and Evolution. Edited by E. Teeling, 37(5), pp. 1530–1534. Available at: 10.1093/molbev/msaa015.

Moody, E.R. et al. (2022) “An estimate of the deepest branches of the tree of life from ancient vertically evolving genes,” eLife, 11, p. e66695. Available at: 10.7554/eLife.66695.

Moran, N.A. and Bennett, G.M. (2014) “The Tiniest Tiny Genomes,” Annual Review of Microbiology, 68(1), pp. 195–215. Available at: 10.1146/annurev-micro-091213-112901.

Moya, A. et al. (2008) “Learning how to live together: genomic insights into prokaryote–animal symbioses,” Nature Reviews Genetics, 9(3), pp. 218–229. Available at: 10.1038/nrg2319.

Nariman, N. et al. (2025) “The Microbiome of an Invasive Spider: Reduced Bacterial Richness, but no Indication of Microbial-Mediated Dispersal Behaviour,” Microbial Ecology, 88(1), p. 70. Available at: 10.1007/s00248-025-02565-6.

Nentwig, W. (1987) “The Prey of Spiders,” in W. Nentwig (ed.) Ecophysiology of Spiders. Berlin, Heidelberg: Springer Berlin Heidelberg, pp. 249–263. Available at: 10.1007/978-3-642-71552-5_18.

Ochman, H. (2005) “Genomes on the shrink,” Proceedings of the National Academy of Sciences, 102(34), pp. 11959–11960. Available at: 10.1073/pnas.0505863102.

O’Fallon, B. (2008) “POPULATION STRUCTURE, LEVELS OF SELECTION, AND THE EVOLUTION OF INTRACELLULAR SYMBIONTS,” Evolution, 62(2), pp. 361–373. Available at: 10.1111/j.1558-5646.2007.00289.x.

Perez-Lamarque, B. et al. (2022) “Limited Evidence for Microbial Transmission in the Phylosymbiosis between Hawaiian Spiders and Their Microbiota,” mSystems. Edited by S.M. Hird, 7(1), pp. e01104–21. Available at: 10.1128/msystems.01104-21.

Pollack, J.D., Williams, M.V. and McElhaney, R.N. (1997) “The Comparative Metabolism of the Mollicutes ( Mycoplasmas): The Utility for Taxonomic Classification and the Relationship of Putative Gene Annotation and Phylogeny to Enzymatic Function in the Smallest Free-Living Cells,” Critical Reviews in Microbiology, 23(4), pp. 269–354. Available at: 10.3109/10408419709115140.

Razin, S. and Hayflick, L. (2010) “Highlights of mycoplasma research—An historical perspective,” Biologicals, 38(2), pp. 183–190. Available at: 10.1016/j.biologicals.2009.11.008.

Razin, S., Yogev, D. and Naot, Y. (1998) “Molecular Biology and Pathogenicity of Mycoplasmas,” Microbiology and Molecular Biology Reviews, 62(4), pp. 1094–1156. Available at: 10.1128/MMBR.62.4.1094-1156.1998.

Rogers, M.J. et al. (1985) “Construction of the mycoplasma evolutionary tree from 5S rRNA sequence data.,” Proceedings of the National Academy of Sciences, 82(4), pp. 1160–1164. Available at: 10.1073/pnas.82.4.1160.

Román-Escrivá, P. et al. (2025) “Metrics of Genomic Complexity in the Evolution of Bacterial Endosymbiosis,” Biology, 14(4), p. 338. Available at: 10.3390/biology14040338.

Seemann, T. (2014) “Prokka: rapid prokaryotic genome annotation,” Bioinformatics, 30(14), pp. 2068–2069. Available at: 10.1093/bioinformatics/btu153.

Seppey, M., Manni, M. and Zdobnov, E.M. (2019) “BUSCO: Assessing Genome Assembly and Annotation Completeness,” in M. Kollmar (ed.) Gene Prediction. New York, NY: Springer New York (Methods in Molecular Biology), pp. 227–245. Available at: 10.1007/978-1-4939-9173-0_14.

Sheffer, M.M. et al. (2020) “Tissue-and Population-Level Microbiome Analysis of the Wasp Spider Argiope bruennichi Identified a Novel Dominant Bacterial Symbiont,” Microorganisms, 8(1), p. 8. Available at: 10.3390/microorganisms8010008.

Sheffer, M.M. et al. (2021) “Chromosome-level reference genome of the European wasp spider Argiope bruennichi□: a resource for studies on range expansion and evolutionary adaptation,” GigaScience, 10(1), p. giaa148. Available at: 10.1093/gigascience/giaa148.

Sheffer, M.M. et al. (2026) “Rapid ecological and evolutionary divergence during a poleward range expansion,” Ecological Monographs, 96(1), p. e70047. Available at: 10.1002/ecm.70047.

Shen, W. et al. (2016) “SeqKit: A Cross-Platform and Ultrafast Toolkit for FASTA/Q File Manipulation,” PLOS ONE. Edited by Q. Zou, 11(10), p. e0163962. Available at: 10.1371/journal.pone.0163962.

Tatusov, R.L. et al. (2003) “The COG database: an updated version includes eukaryotes,” BMC Bioinformatics, 4(1), p. 41. Available at: 10.1186/1471-2105-4-41.

Tatusova, T. et al. (2016) “NCBI prokaryotic genome annotation pipeline,” Nucleic Acids Research, 44(14), pp. 6614–6624. Available at: 10.1093/nar/gkw569.

The UniProt Consortium (2019) “UniProt: a worldwide hub of protein knowledge,” Nucleic Acids Research, 47(D1), pp. D506–D515. Available at: 10.1093/nar/gky1049.

Toft, C. and Andersson, S.G.E. (2010) “Evolutionary microbial genomics: insights into bacterial host adaptation,” Nature Reviews Genetics, 11(7), pp. 465–475. Available at: 10.1038/nrg2798.

Vanthournout, B. and Hendrickx, F. (2015) “Endosymbiont Dominated Bacterial Communities in a Dwarf Spider,” PLOS ONE. Edited by M. Horn, 10(2), p. e0117297. Available at: 10.1371/journal.pone.0117297.

Vaser, R. et al. (2017) “Fast and accurate de novo genome assembly from long uncorrected reads,” Genome Research, 27(5), pp. 737–746. Available at: 10.1101/gr.214270.116.

Vaser, R. and Šikić, M. (2019) “Yet another de novo genome assembler.” Bioinformatics. Available at: 10.1101/656306.

Walker, B.J. et al. (2014) “Pilon: An Integrated Tool for Comprehensive Microbial Variant Detection and Genome Assembly Improvement,” PLoS ONE. Edited by J. Wang, 9(11), p. e112963. Available at: 10.1371/journal.pone.0112963.

Wang, Y. and Chandler, C. (2016) “Candidate pathogenicity islands in the genome of ‘ Candidatus Rickettsiella isopodorum’, an intracellular bacterium infecting terrestrial isopod crustaceans,” PeerJ, 4, p. e2806. Available at: 10.7717/peerj.2806.

Wang, Y.-Y. et al. (2026) “The reduced genome of Candidatus Portiera sp. in Bemisia afer: evolutionary trajectories and functional implications,” BMC Genomics, 27(1), p. 205. Available at: 10.1186/s12864-025-12509-6.

Wawer, W. et al. (2017) “Population structure of the expansive wasp spider ( Argiope bruennichi) at the edge of its range,” Journal of Arachnology, 45(3), pp. 361–369. Available at: 10.1636/JoA-S-16-056.1.

Wawer, W. and Wytwer, J. (2020) “Abundance changes in orb-weaver spider communities at the edge of the Argiope bruennichi expansion range,” Zootaxa, 4899(1). Available at: 10.11646/zootaxa.4899.1.18.

White, J.A. et al. (2020) “Endosymbiotic Bacteria Are Prevalent and Diverse in Agricultural Spiders,” Microbial Ecology, 79(2), pp. 472–481. Available at: 10.1007/s00248-019-01411-w.

Wick, R.R. et al. (2015) “Bandage: interactive visualization of de novo genome assemblies,” Bioinformatics, 31(20), pp. 3350–3352. Available at: 10.1093/bioinformatics/btv383.

Wickham, H. et al. (2019) “Welcome to the Tidyverse,” Journal of Open Source Software, 4(43), p. 1686. Available at: 10.21105/joss.01686.

Wolz, M. et al. (2020) “Dispersal and life-history traits in a spider with rapid range expansion,” Movement Ecology, 8(1), p. 2. Available at: 10.1186/s40462-019-0182-4.

Yavlovich, A., Tarshis, M. and Rottem, S. (2004) “Internalization and intracellular survival of Mycoplasma pneumoniae by non-phagocytic cells,” FEMS Microbiology Letters, 233(2), pp. 241–246. Available at: 10.1111/j.1574-6968.2004.tb09488.x.

Zhang, L. et al. (2017) “Bacterial community of a spider, Marpiss magister (Salticidae),” 3 Biotech, 7(6), p. 371. Available at: 10.1007/s13205-017-0994-0.

Zhang, L. et al. (2018) “Insights into the bacterial symbiont diversity in spiders,” Ecology and Evolution, 8(10), pp. 4899–4906. Available at: 10.1002/ece3.4051.

