## Supplementary figures and images for "Genomic insights into *Candidatus* Argioplasma dusa, a bacterial symbiont of the wasp spider *Argiope bruennichi*"

### Fig. S1

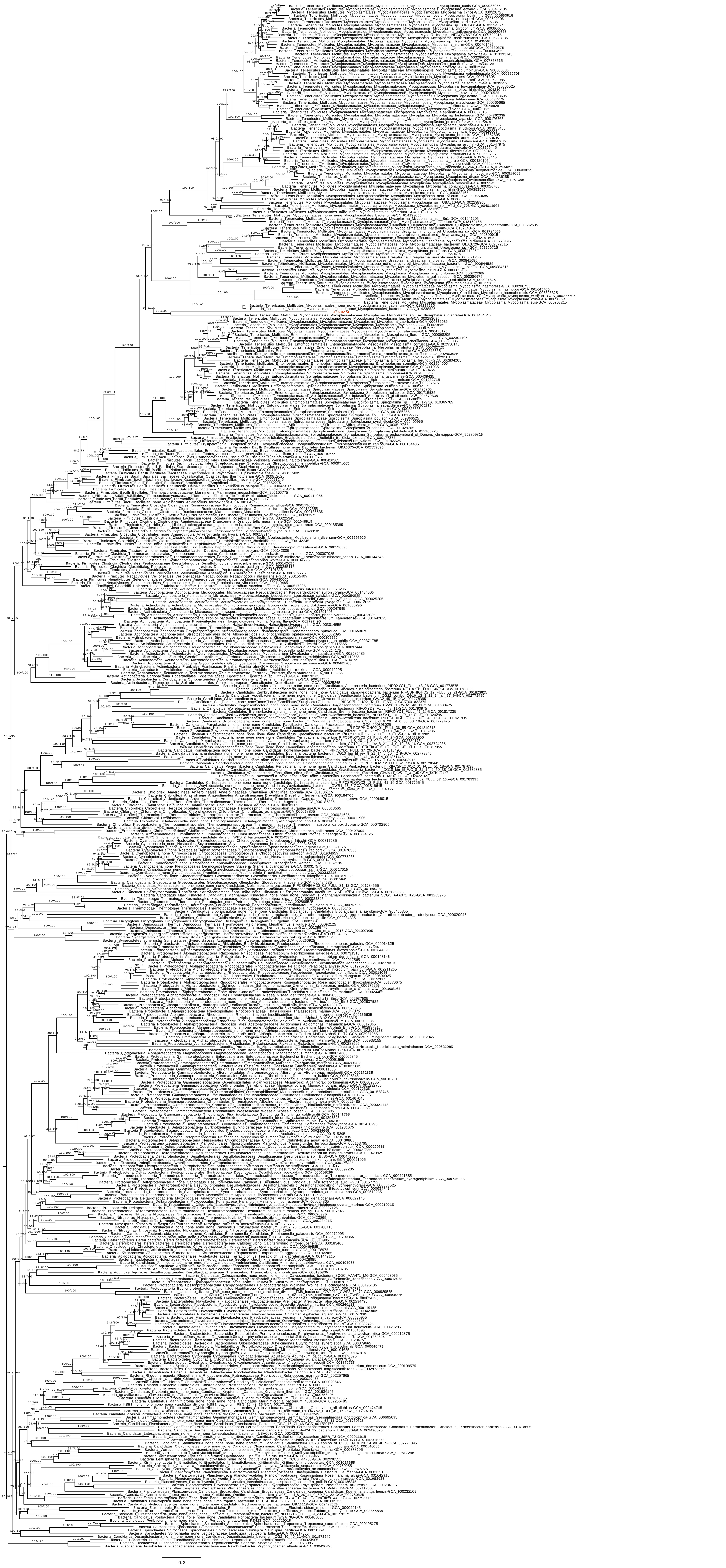

### Fig. S2

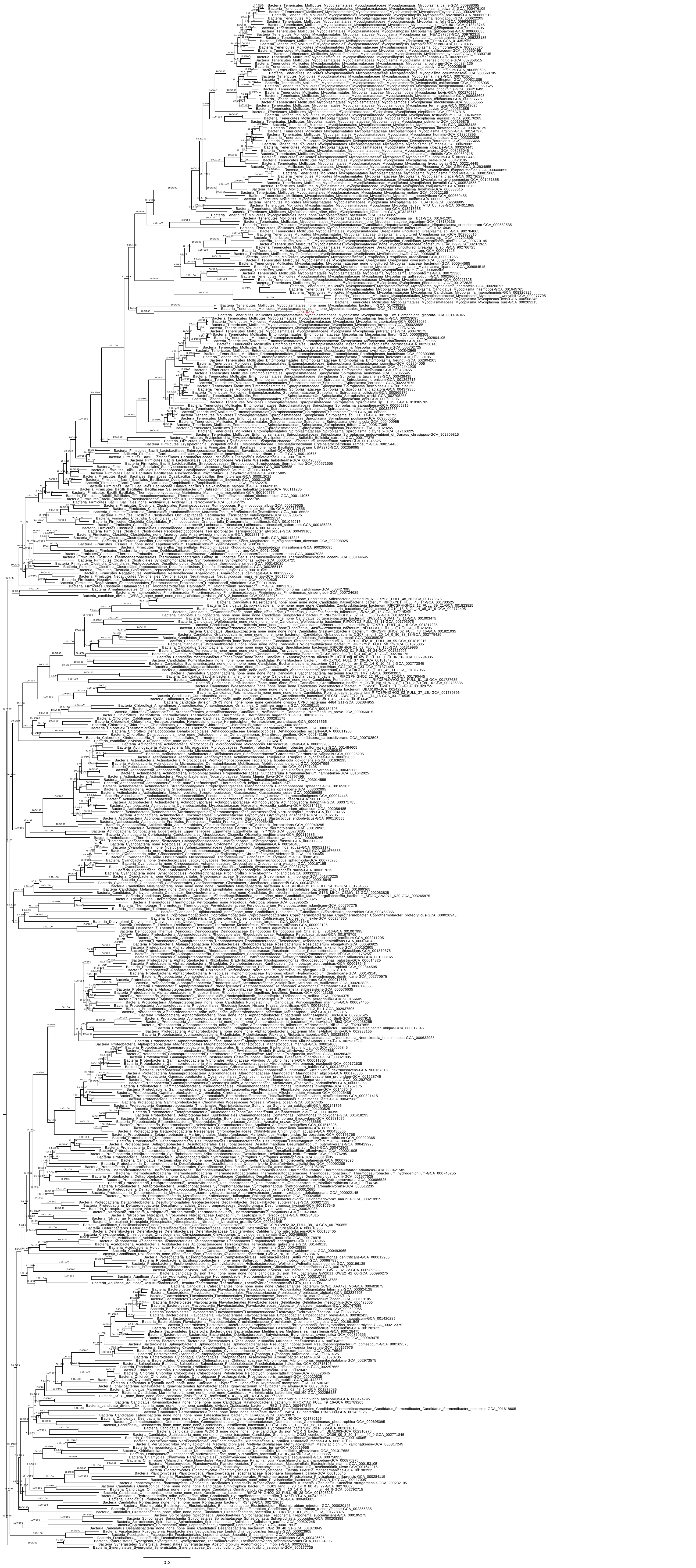

### Fig. S3

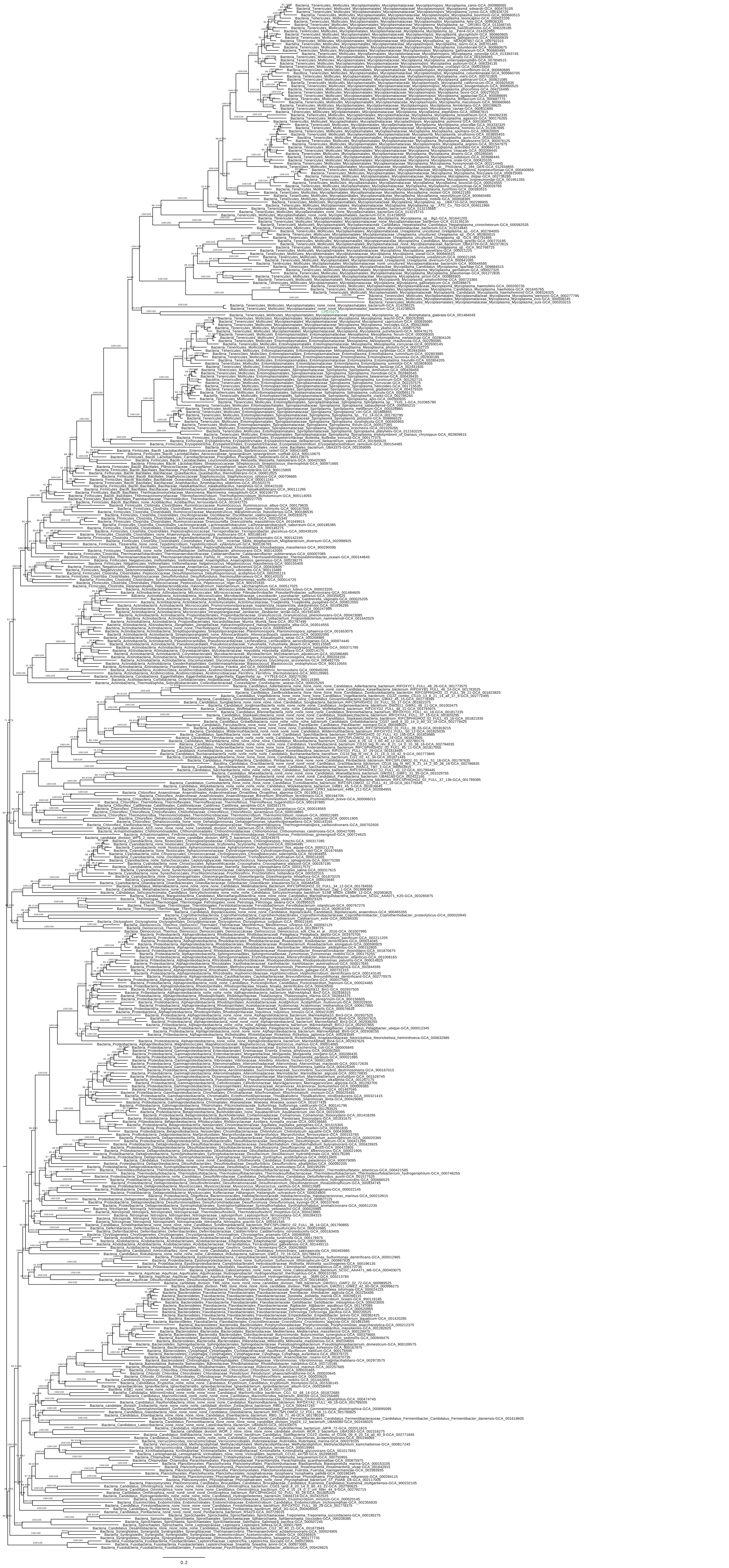

### Fig. S4

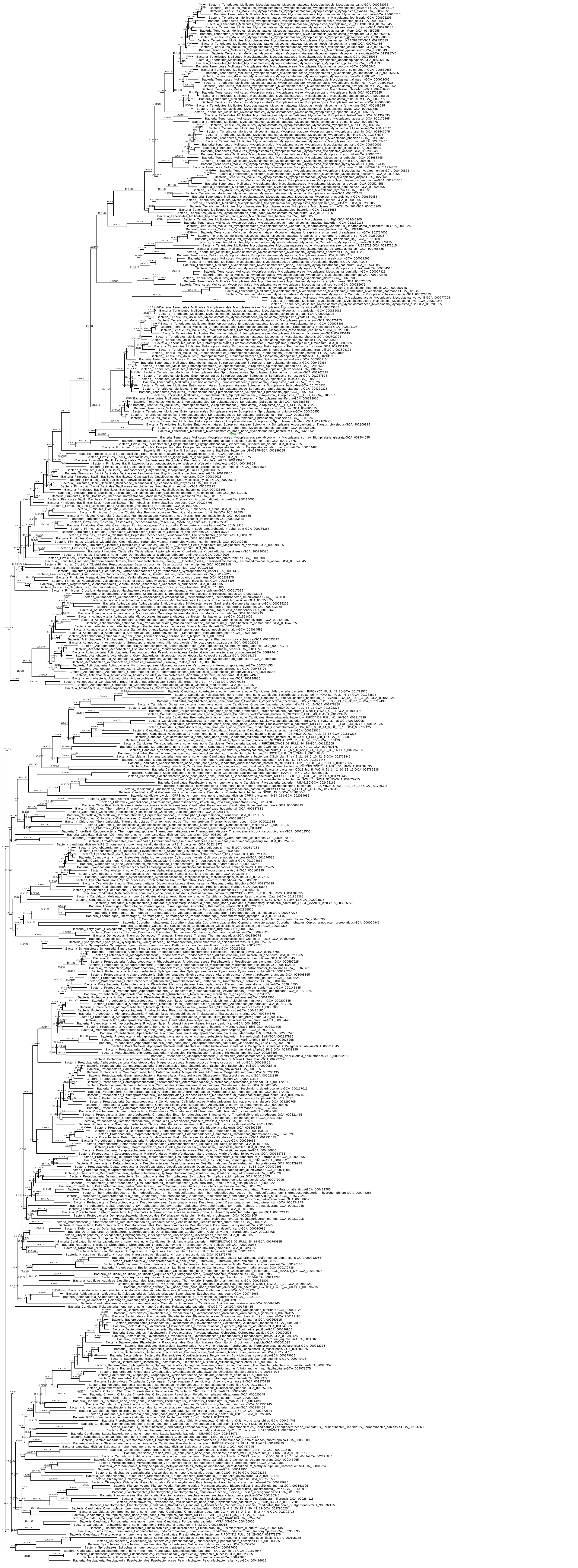

### Fig. S5

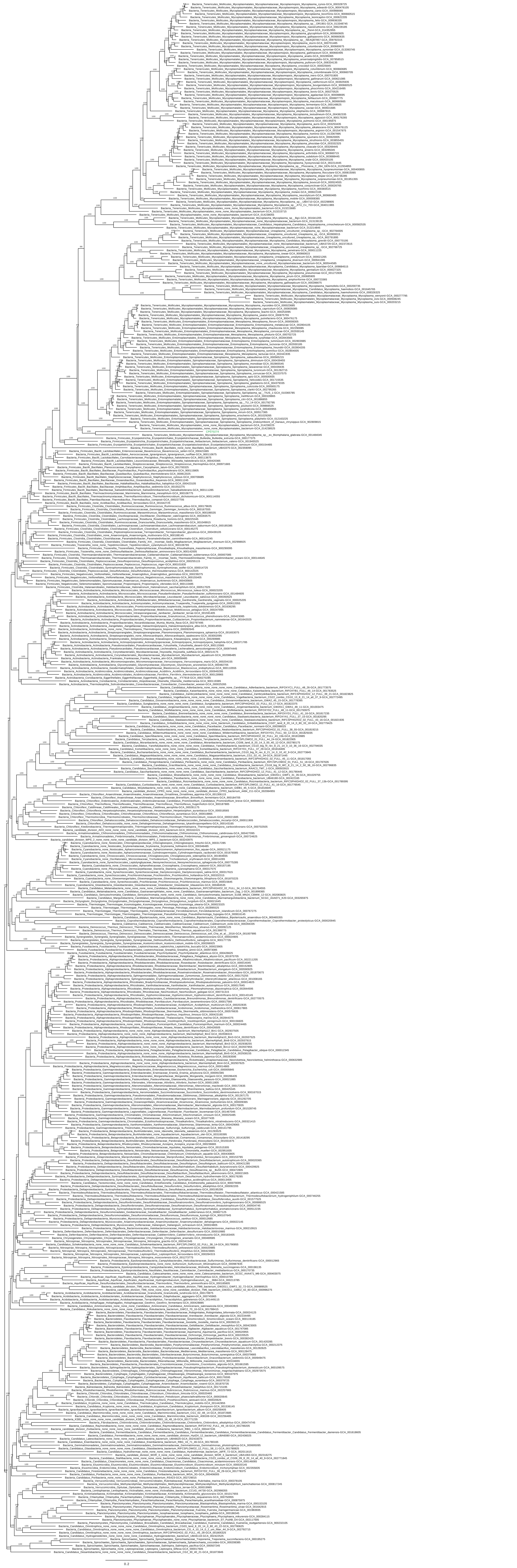

### Fig. S6

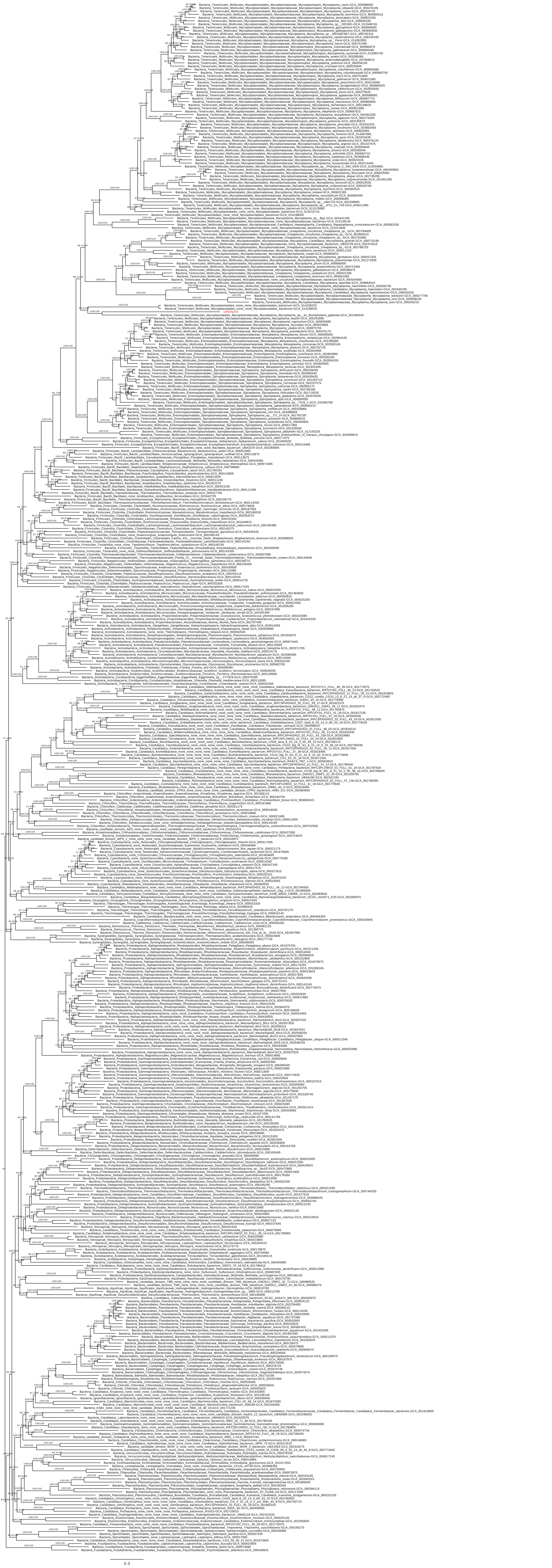

### Fig. S7

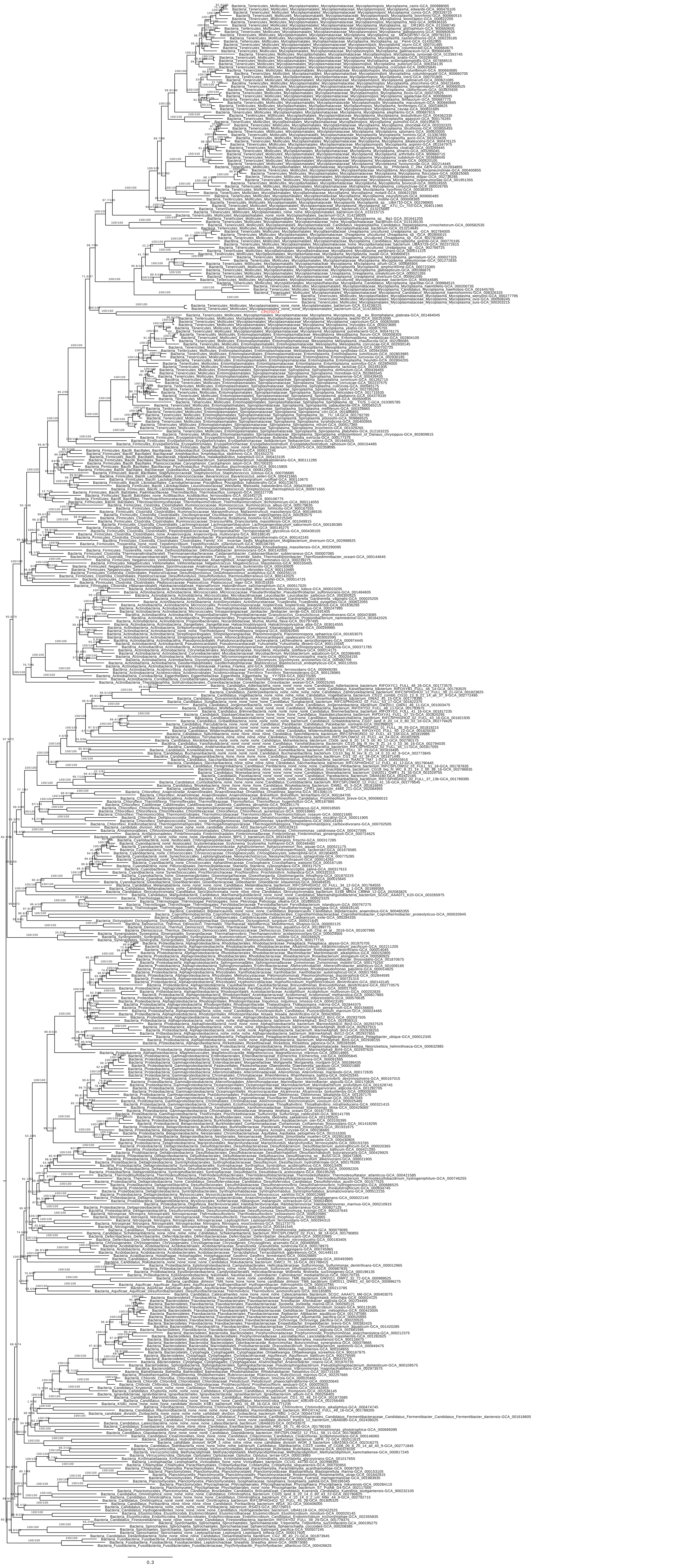

### Fig. S8

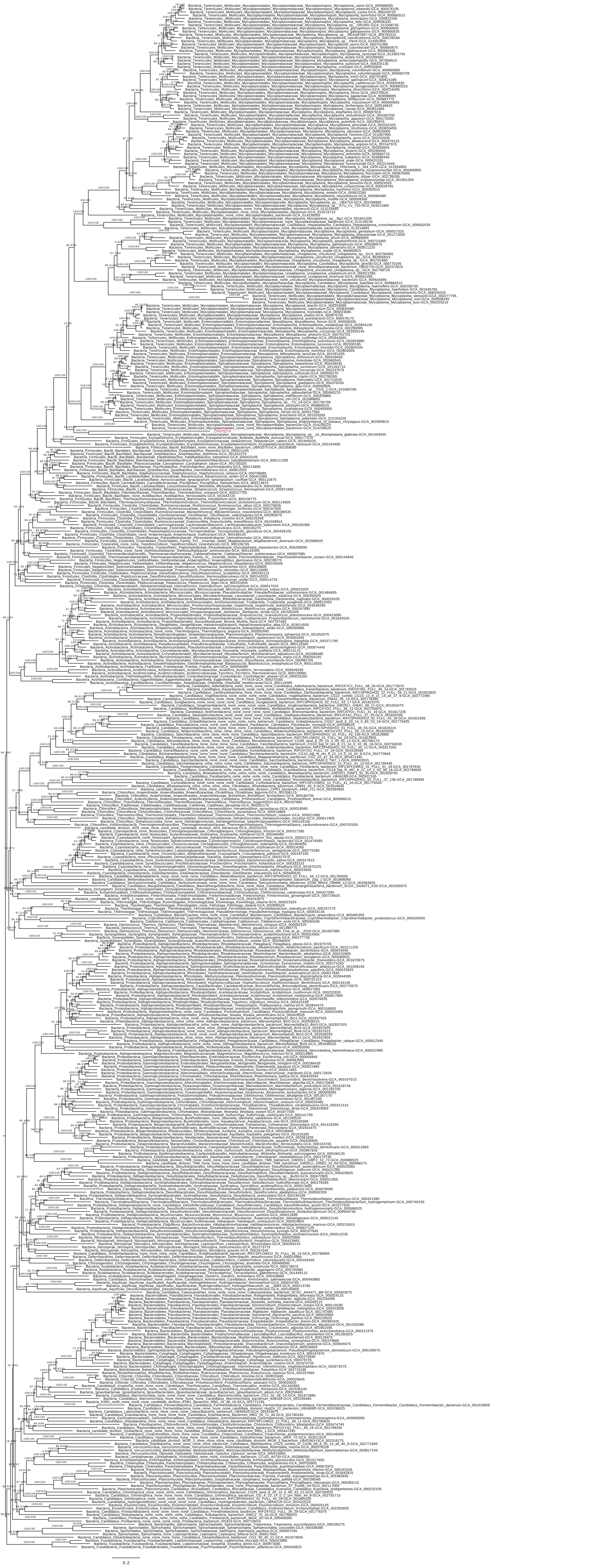

### Fig. S9

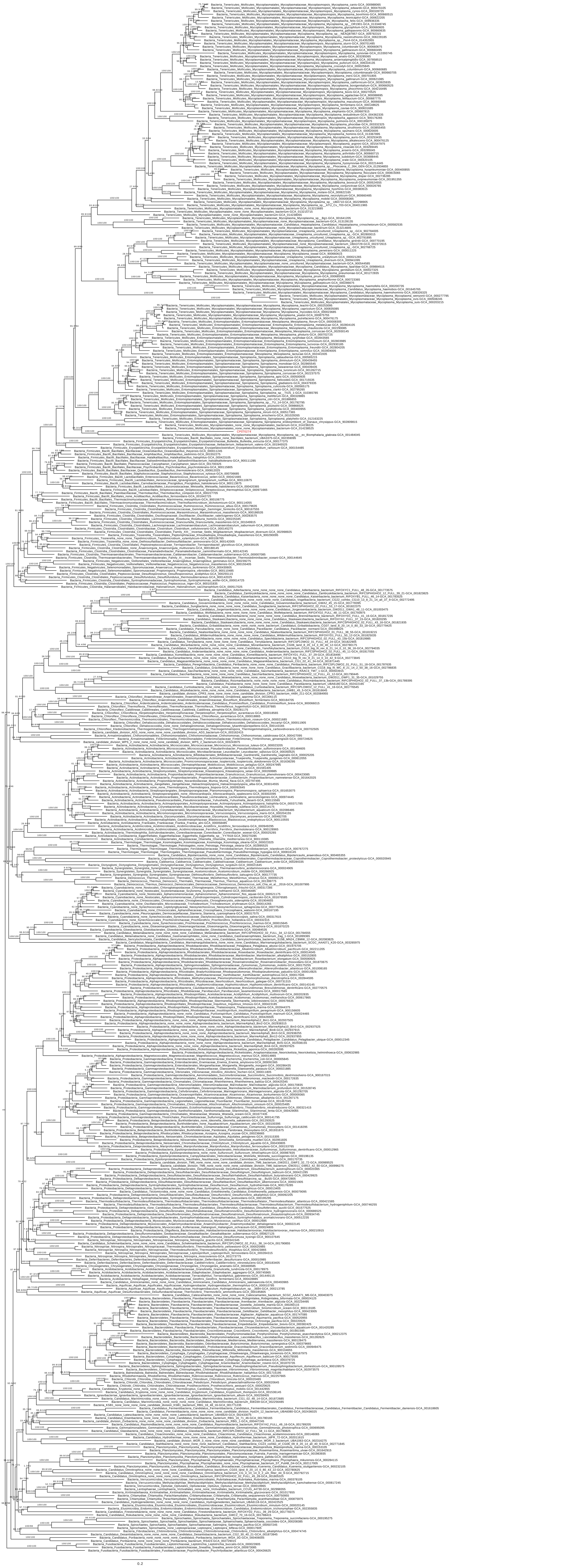
