## Supplemental Data for "Genomic insights into *Candidatus* Argioplasma dusa, a bacterial symbiont of the wasp spider *Argiope bruennichi*"

^6^Current address: Department of Biological Sciences, California Polytechnic University, Humboldt, Arcata, CA, 95521 USA

*shared first authorship

^‡^corresponding author

**Supplementary Methods**

**Method S1: Assembly of the *Candidatus* Argioplasma dusa genome**

**Read preparation**

**PacBio reads** (Sequence Read Archive: SRR11652932) (1) were quality (top 90%) and length (>1000 bp) trimmed using Filtlong and duplication cleaned by the SeqKit (2) rmdup function. After trimming, 3978330 PacBio reads (∼68% of the raw reads) with a mean length of 7,512.13 bp and a length range of 1000 bp to 101,537 bp remained.

**Illumina paired-end reads** (European Nucleotide Archive: ERR574428) (3) were used for assembly polishing. Illumina reads were adapter, quality and length trimmed by Trimmomatic (4). The default parameters were chosen. The provided TruSeq3-PE adapter sequences (PrefixPE/1:TACACTCTTTCCCTACACGACGCTCTTCCGATCT; PrefixPE/2:GTGACTGGAGTTCAGACGTGTGCTCTTCCGATCT) were used for adapter trimming. We *in silico* pooled data of five populations from Portugal, Italy, Sweden, Baltic region and Japan. In total, 843,533,777 paired sequences with a length range of 36 bp to 101 bp remained. The mean forward read length was 99.91 bp and the average reverse read length was 99.49 bp.

**Assembly strategies**

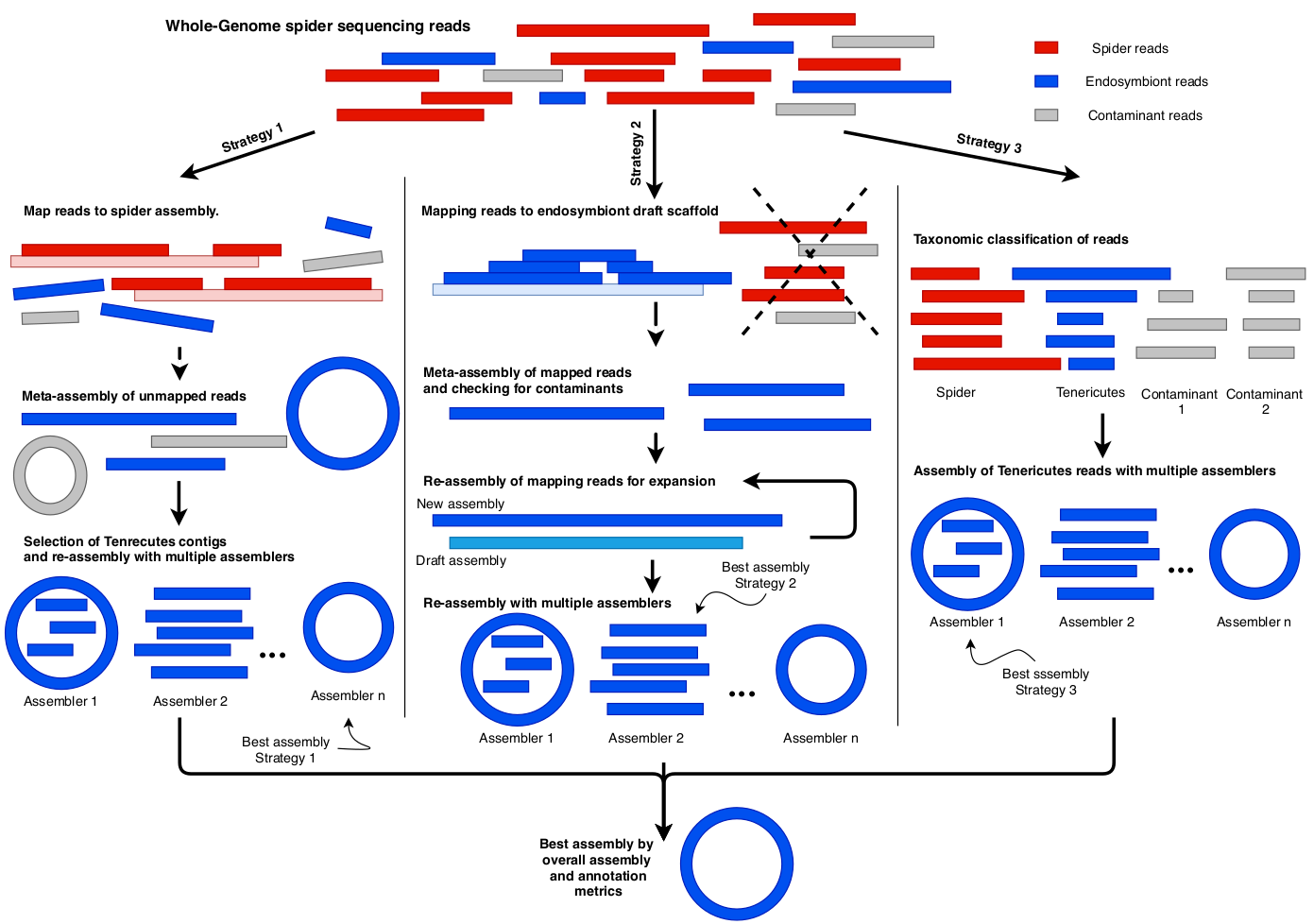

**Fig. S10: Overview of assembly strategies.** Strategy 1 aimed to select whole genome spider reads that do not map to the spider assembly and to assemble those reads. Strategy 2 aimed to expand and improve a Miniasm (5) endosymbiont draft scaffold. Strategy 3 aimed to classify whole genome spider reads and to assemble Tenericutes reads.

**Strategy 1**

Whole genome *A. bruennichi* spider reads were aligned against the *A. bruennichi* genome (1) using Minimap2 (6) for trimmed PacBio reads and BWA-MEM (7) for trimmed Illumina reads. Reads, which do not map to the spider genome assembly, were filtered using SAMtools (8). Resulting singleton Illumina reads were discarded. PacBio reads were assembled as metagenomic data using metaFlye (9). The resulting meta-assembly was polished twice with long reads using Racon (10) and twice with short reads using Pilon (11). The polished meta-assembly was investigated using BlobTools (12) and DIAMOND (13) with the UniProt (14) database for taxonomic classification. We identified two Tenericutes contigs comprising to about 120 kb. We then performed the “map-and-assemble” approach (15), meaning to iteratively map all reads on the selected template and to reassemble mapping reads, on the two Tenericutes contigs. In addition, we also performed the “map-and-assemble” approach on the two Tenericutes contigs, all Firmicutes contigs (since Tenericutes belong to Firmicutes) and no-hit contigs in a GC content range of 20% to 40% (Tenericutes genomes are ~30% GC (16)). For the two Tenericutes contigs, five iterations were performed to gain a circular draft assembly containing a sequence with at least 90% identity to the previous identified 16S rDNA sequence of *Ca.* A. dusa (17) using Flye (18). For the Tenericutes, Firmicutes and no-hit contigs, the assembly was reassembled in three iterations using metaFlye and processing with BlobTools to select Tenericutes, Firmicutes and no-hit contigs. Assembly contigs were investigated using Bandage (19) and QUAST (20).

**Strategy 2**

Reads were mapped onto a 587,333 bp symbiont scaffold generated as contaminant contig in the baseline assembly in the *A. bruennichi* assembly process (1) using Minimap2 and BWA-MEM. This contig was associated to *Ca.* A. dusa by identifying the previous generated 16S rDNA sequence (17). Mapping reads were selected using SAMtools and assembled as metagenomic data using metaFlye. The resulting assembly was polished twice with long reads using Racon and twice with short reads using Pilon. Tenericutes contigs identified by BlobTools were selected and reassembled by performing the “map-and-assemble” approach in four iterations using Flye. Contigs were investigated using Bandage and QUAST. The Tenericutes contig was selected as draft assembly.

**Strategy 3**

PacBio reads were binned using BlobTools/DIAMOND. In this case, we treated the long PacBio reads as small assembly contigs. For this, reads were converted from FastQ to FASTA format with seqkit fq2fa. Then, reads classified as Tenericutes were selected and assembled with Flye. This resulted in a fragmented assembly. However, as a second approach, we mapped all PacBio reads to the Tenericutes-classified PacBio reads using Minimap2 and the mapped reads were assembled with Flye (with default polishing step). The assemblies were checked with Bandage for circularity and for *Ca.* A. dusa 16S rRNA sequence presence using Blast+. The resulting assembly was considered a draft assembly.

**Final assembling**

PacBio reads were mapped onto the draft assemblies and mapped reads were assembled with Flye, Canu (21), Raven (22), Redbean (23) and miniasm+minipolish (24) (each with its default polishing steps). Assemblies were checked with Bandage for circularity and *Ca.* A. dusa 16S rRNA sequence presence. Resulting assemblies were then polished twice with Racon using PacBio reads and twice with Pilon using Illumina reads. Polished assemblies were checked with BlobTools for contaminants and compared with QUAST for basic assembly metrics, with Mauve (25) for structural similarity and with BUSCO (26) for conserved Tenericutes genes.
The final assembly coming from strategy 3 with reassembly using Raven was chosen based on BUSCO completeness using the OrthoDB Tenericutes reference and largest one-contig assembly.

### **Method S2: Functional Annotation**

For further functional annotation, concatenated proteins from all *Ca*. A. dusa and the 478 bacterial reference genomes were compared against several databases, including the COGs (using the 2020 update)

(27), arCOGs (version from 2020) (28), the KO profiles from the KEGG Automatic Annotation Server (KAAS; downloaded April 2019) (29), the Pfam database (Release 31.0) (30), the TIGRFAM database (Release 15.0) (31), the CAZymes database(dbCAN-HMMdb-V7) (32), the hydrogenase database (downloaded in November, 2018) (33) and NCBIs non-redundant (nr) database (downloaded in 2019) (34). Individual database searches, with the exception of searches against the hydrogenase and nr database, were conducted by querying each individual database against all concatenated proteins using hmmsearch (v3.1b2; settings: --tblout --domtblout --notextw) (35). The output was parsed to only include hits with an e-value less than 1e-3 and only a single hit for each protein was selected based on the best e-value and bit score value. The hydrogenase database was queried using BLASTp (v2.7.1+, settings: -outfmt 6 -evalue 1e-10) (36) and the nr database was queried with DIAMOND (v0.9.22.23, settings: diamond blastp --more-sensitive --evalue 1e-3 --seq 50 --db $ncbi_nr_db --taxonmap $taxon_db --outfmt 6 qseqid qtitle qlen sseqid salltitles slen qstart qend sstart send evalue bitscore length pident staxids) (13). Additionally, all proteins were scanned for protein domains using InterProScan (v5.29-68.0; settings: --iprlookup --goterms) (37).

**Supplementary Data**

| **BuscoID** | **Description** | **% in Tenericutes** |
| --- | --- | --- |
| 10044at544448 | Ribosomal protein L35 | 91.9 |
| 1484at544448 | DNA polymerase III subunit gamma/tau | 99.2 |
| 2088at544448 | GTPase Der | 100.0 |
| 2803at544448 | Pseudouridine synthase | 98.4 |
| 2862at544448 | DhaL domain | 90.3 |
| 3750at544448 | tRNA modification GTPase MnmE | 96.0 |
| 4126at544448 | Energy-coupling factor transporter ATP-binding protein EcfA | 100.0 |
| 4491at544448 | Ribosomal protein L1 | 98.4 |
| 4797at544448 | Cytidylate kinase | 96.0 |
| 4972at544448 | GTPase Era | 94.4 |
| 5381at544448 | tRNA N6-adenosine threonylcarbamoyltransferase | 98.4 |
| 5957at544448 | Chromosomal replication initiator protein DnaA | 94.4 |
| 6113at544448 | rRNA (cytidine-2’-O-)-methyltransferase | 93.5 |
| 6553at544448 | RNA 2-O ribose methyltransferase, substrate binding | 96.0 |
| 6666at544448 | Thymidine kinase | 95.2 |
| 7157at544448 | Ribosome biogenesis GTPase RsgA | 92.7 |
| 7177at544448 | Uracil phosphoribosyltransferase | 91.9 |
| 723at544448 | Methionine–tRNA ligase | 99.2 |
| 7503at544448 | Holliday junction resolvase RecU | 89.5 |
| 7514at544448 | tRNA pseudouridine synthase B | 91.1 |
| 7555at544448 | Ribosomal protein S8 | 94.4 |
| 8206at544448 | Ribonuclease P | 90.3 |
| 8505at544448 | Gcp-like domain | 90.3 |
| 9641at544448 | Ribosomal protein L7/L12, C-terminal | 99.2 |

**Table S2:** List of missing Tenericutes BUSCOs. After annotation of the generated assembly with PGAP the BUSCO completeness was determined according to OrthoDB v10.1 Tenericutes orthologues database. 24 BUSCOs were found missing. The missing BUSCOs are presented with their description and the percentage of appearance in the 124 Tenericutes reference species.

| **Single specimen/ pooled** | **sample** | **# reads** | **# mapped reads** | **% mapped reads** |
| --- | --- | --- | --- | --- |
| single | Japan20 | 72390374 | 185436 | 0.256 |
| single | Japan221 | 91479354 | 3510 | 0.004 |
| single | Japan251 | 70256859 | 427519 | 0.609 |
| single | Japan271 | 79726374 | 164 | 0.000 |
| single | Japan281 | 67381722 | 134 | 0.000 |
| pooled | Sweden | 355912405 | 1897732 | 0.533 |
| pooled | Baltic | 321871789 | 1866854 | 0.580 |
| pooled | Portugal | 342637709 | 2177960 | 0.636 |
| pooled | Italy | 285414033 | 1465467 | 0.513 |
| pooled | Azores | 332886877 | 1411863 | 0.424 |
| single | Madeira | 356322474 | 522058 | 0.147 |

**Table S3:** Illumina reads generated from different *A. bruennichi* populations.

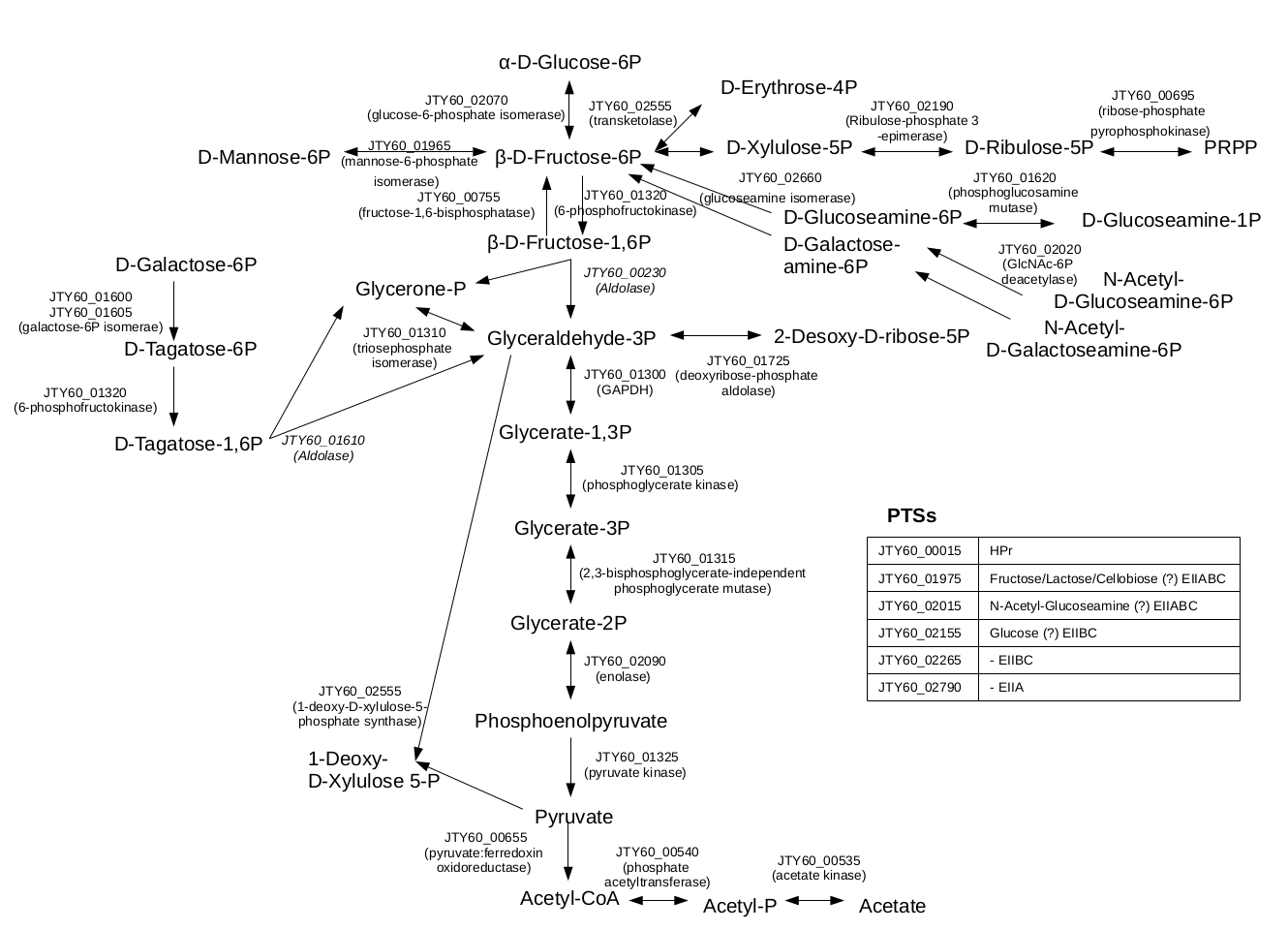

**Figure S11:** Detailed schematic visualization of the central C-metabolism of *Ca.* A. dusa

**References**

1.	Sheffer MM, Hoppe A, Krehenwinkel H, Uhl G, Kuss AW, Jensen L, Jensen C, Gillespie RG, Hoff KJ, Prost S. 2021. Chromosome-level reference genome of the European wasp spider *Argiope bruennichi* : a resource for studies on range expansion and evolutionary adaptation. GigaScience 10:giaa148. https://doi.org/10.1093/gigascience/giaa148.

2.	Shen W, Le S, Li Y, Hu F. 2016. SeqKit: A Cross-Platform and Ultrafast Toolkit for FASTA/Q File Manipulation. PLoS ONE 11:e0163962. https://doi.org/10.1371/journal.pone.0163962.

3.	Krehenwinkel H, Rödder D, Tautz D. 2015. Eco‐genomic analysis of the poleward range expansion of the wasp spider *A rgiope bruennichi* shows rapid adaptation and genomic admixture. Global Change Biology 21:4320–4332. https://doi.org/10.1111/gcb.13042.

4.	Bolger AM, Lohse M, Usadel B. 2014. Trimmomatic: a flexible trimmer for Illumina sequence data. Bioinformatics 30:2114–2120. https://doi.org/10.1093/bioinformatics/btu170.

5.	Li H. 2016. Minimap and miniasm: fast mapping and de novo assembly for noisy long sequences. Bioinformatics 32:2103–2110. https://doi.org/10.1093/bioinformatics/btw152.

6.	Li H. 2018. Minimap2: pairwise alignment for nucleotide sequences. Bioinformatics 34:3094–3100. https://doi.org/10.1093/bioinformatics/bty191.

7.	Li H. 2013. Aligning sequence reads, clone sequences and assembly contigs with BWA-MEM (2). arXiv https://doi.org/10.48550/ARXIV.1303.3997. https://doi.org/10.48550/ARXIV.1303.3997.

8.	Li H, Handsaker B, Wysoker A, Fennell T, Ruan J, Homer N, Marth G, Abecasis G, Durbin R, 1000 Genome Project Data Processing Subgroup. 2009. The Sequence Alignment/Map format and SAMtools. Bioinformatics 25:2078–2079. https://doi.org/10.1093/bioinformatics/btp352.

9.	Kolmogorov M, Bickhart DM, Behsaz B, Gurevich A, Rayko M, Shin SB, Kuhn K, Yuan J, Polevikov E, Smith TPL, Pevzner PA. 2020. metaFlye: scalable long-read metagenome assembly using repeat graphs. Nat Methods 17:1103–1110. https://doi.org/10.1038/s41592-020-00971-x.

10.	Vaser R, Sović I, Nagarajan N, Šikić M. 2017. Fast and accurate de novo genome assembly from long uncorrected reads. Genome Res 27:737–746. https://doi.org/10.1101/gr.214270.116.

11.	Walker BJ, Abeel T, Shea T, Priest M, Abouelliel A, Sakthikumar S, Cuomo CA, Zeng Q, Wortman J, Young SK, Earl AM. 2014. Pilon: An Integrated Tool for Comprehensive Microbial Variant Detection and Genome Assembly Improvement. PLoS ONE 9:e112963. https://doi.org/10.1371/journal.pone.0112963.

12.	Laetsch DR, Blaxter ML. 2017. BlobTools: Interrogation of genome assemblies. F1000Res 6:1287. https://doi.org/10.12688/f1000research.12232.1.

13.	Buchfink B, Xie C, Huson DH. 2015. Fast and sensitive protein alignment using DIAMOND. Nat Methods 12:59–60. https://doi.org/10.1038/nmeth.3176.

14.	The UniProt Consortium. 2019. UniProt: a worldwide hub of protein knowledge. Nucleic Acids Research 47:D506–D515. https://doi.org/10.1093/nar/gky1049.

15.	Wang Y, Chandler C. 2016. Candidate pathogenicity islands in the genome of ‘ *Candidatus* Rickettsiella isopodorum’, an intracellular bacterium infecting terrestrial isopod crustaceans. PeerJ 4:e2806. https://doi.org/10.7717/peerj.2806.

16.	Reichenberger ER, Rosen G, Hershberg U, Hershberg R. 2015. Prokaryotic Nucleotide Composition Is Shaped by Both Phylogeny and the Environment. Genome Biology and Evolution 7:1380–1389. https://doi.org/10.1093/gbe/evv063.

17.	Sheffer MM, Uhl G, Prost S, Lueders T, Urich T, Bengtsson MM. 2019. Tissue- and Population-Level Microbiome Analysis of the Wasp Spider Argiope bruennichi Identified a Novel Dominant Bacterial Symbiont. Microorganisms 8:8. https://doi.org/10.3390/microorganisms8010008.

18.	Kolmogorov M, Yuan J, Lin Y, Pevzner PA. 2019. Assembly of long, error-prone reads using repeat graphs. Nat Biotechnol 37:540–546. https://doi.org/10.1038/s41587-019-0072-8.

19.	Wick RR, Schultz MB, Zobel J, Holt KE. 2015. Bandage: interactive visualization of *de novo* genome assemblies. Bioinformatics 31:3350–3352. https://doi.org/10.1093/bioinformatics/btv383.

20.	Gurevich A, Saveliev V, Vyahhi N, Tesler G. 2013. QUAST: quality assessment tool for genome assemblies. Bioinformatics 29:1072–1075. https://doi.org/10.1093/bioinformatics/btt086.

21.	Koren S, Walenz BP, Berlin K, Miller JR, Bergman NH, Phillippy AM. 2017. Canu: scalable and accurate long-read assembly via adaptive *k* -mer weighting and repeat separation. Genome Res 27:722–736. https://doi.org/10.1101/gr.215087.116.

22.	Vaser R, Šikić M. 2019. Yet another de novo genome assembler. Bioinformatics https://doi.org/10.1101/656306. https://doi.org/10.1101/656306.

23.	Ruan J, Li H. 2020. Fast and accurate long-read assembly with wtdbg2. Nat Methods 17:155–158. https://doi.org/10.1038/s41592-019-0669-3.

24.	Wick RR, Holt KE. 2021. Benchmarking of long-read assemblers for prokaryote whole genome sequencing. F1000Res 8:2138. https://doi.org/10.12688/f1000research.21782.4.

25.	Darling ACE, Mau B, Blattner FR, Perna NT. 2004. Mauve: Multiple Alignment of Conserved Genomic Sequence With Rearrangements. Genome Res 14:1394–1403. https://doi.org/10.1101/gr.2289704.

26.	Seppey M, Manni M, Zdobnov EM. 2019. BUSCO: Assessing Genome Assembly and Annotation Completeness, p. 227–245. *In* Kollmar, M (ed.), Gene Prediction. Springer New York, New York, NY. https://doi.org/10.1007/978-1-4939-9173-0_14.

27.	Tatusov RL, Fedorova ND, Jackson JD, Jacobs AR, Kiryutin B, Koonin EV, Krylov DM, Mazumder R, Mekhedov SL, Nikolskaya AN, Rao BS, Smirnov S, Sverdlov AV, Vasudevan S, Wolf YI, Yin JJ, Natale DA. 2003. The COG database: an updated version includes eukaryotes. BMC Bioinformatics 4:41. https://doi.org/10.1186/1471-2105-4-41.

28.	Makarova K, Wolf Y, Koonin E. 2015. Archaeal Clusters of Orthologous Genes (arCOGs): An Update and Application for Analysis of Shared Features between Thermococcales, Methanococcales, and Methanobacteriales. Life 5:818–840. https://doi.org/10.3390/life5010818.

29.	Aramaki T, Blanc-Mathieu R, Endo H, Ohkubo K, Kanehisa M, Goto S, Ogata H. 2020. KofamKOALA: KEGG Ortholog assignment based on profile HMM and adaptive score threshold. Bioinformatics 36:2251–2252. https://doi.org/10.1093/bioinformatics/btz859.

30.	Bateman A. 2004. The Pfam protein families database. Nucleic Acids Research 32:138D – 141. https://doi.org/10.1093/nar/gkh121.

31.	Haft DH. 2003. The TIGRFAMs database of protein families. Nucleic Acids Research 31:371–373. https://doi.org/10.1093/nar/gkg128.

32.	Yin Y, Mao X, Yang J, Chen X, Mao F, Xu Y. 2012. dbCAN: a web resource for automated carbohydrate-active enzyme annotation. Nucleic Acids Research 40:W445–W451. https://doi.org/10.1093/nar/gks479.

33.	Søndergaard D, Pedersen CNS, Greening C. 2016. HydDB: A web tool for hydrogenase classification and analysis. Sci Rep 6:34212. https://doi.org/10.1038/srep34212.

34.	O’Leary NA, Wright MW, Brister JR, Ciufo S, Haddad D, McVeigh R, Rajput B, Robbertse B, Smith-White B, Ako-Adjei D, Astashyn A, Badretdin A, Bao Y, Blinkova O, Brover V, Chetvernin V, Choi J, Cox E, Ermolaeva O, Farrell CM, Goldfarb T, Gupta T, Haft D, Hatcher E, Hlavina W, Joardar VS, Kodali VK, Li W, Maglott D, Masterson P, McGarvey KM, Murphy MR, O’Neill K, Pujar S, Rangwala SH, Rausch D, Riddick LD, Schoch C, Shkeda A, Storz SS, Sun H, Thibaud-Nissen F, Tolstoy I, Tully RE, Vatsan AR, Wallin C, Webb D, Wu W, Landrum MJ, Kimchi A, Tatusova T, DiCuccio M, Kitts P, Murphy TD, Pruitt KD. 2016. Reference sequence (RefSeq) database at NCBI: current status, taxonomic expansion, and functional annotation. Nucleic Acids Res 44:D733–D745. https://doi.org/10.1093/nar/gkv1189.

35.	Finn RD, Clements J, Eddy SR. 2011. HMMER web server: interactive sequence similarity searching. Nucleic Acids Research 39:W29–W37. https://doi.org/10.1093/nar/gkr367.

36.	Altschul SF, Gish W, Miller W, Myers EW, Lipman DJ. 1990. Basic local alignment search tool. Journal of Molecular Biology 215:403–410. https://doi.org/10.1016/S0022-2836(05)80360-2.

37.	Jones P, Binns D, Chang H-Y, Fraser M, Li W, McAnulla C, McWilliam H, Maslen J, Mitchell A, Nuka G, Pesseat S, Quinn AF, Sangrador-Vegas A, Scheremetjew M, Yong S-Y, Lopez R, Hunter S. 2014. InterProScan 5: genome-scale protein function classification. Bioinformatics 30:1236–1240. https://doi.org/10.1093/bioinformatics/btu031.
